# DynaFold: A Latent Diffusion Based Generative Framework for Protein Dynamic Trajectory

**DOI:** 10.1101/2025.09.14.676071

**Authors:** Zirui Fan, Junjie Zhu, Hai-Feng Chen

## Abstract

The dynamic process of protein folding and conformation switching describes the basis of protein functions. Molecular dynamics (MD) simulations are precise computational tools for exploring protein dynamics, but the high computational costs make it difficult to scale up. Deep learning methods have been used to model the Boltzmann distribution of molecular simulations, but achieving MD-level accuracy remains a major challenge. Here, we present DynaFold, a generative deep learning framework based on latent diffusion for sampling protein dynamic trajectories. DynaFold accepts an initial structure and generalizes the conformational dynamics of different proteins with minimal trajectory data during training. It achieves state-of-the-art accuracy in predicting conformational ensembles and sampling conformational transition pathways, demonstrating superior generalization capability and computational efficiency compared to existing methods. Our framework provides a general solution for generating conformation distributions and transition processes between different conformations for proteins, enabling rapid sampling of structural ensembles and analysis of Boltzmann systems.

## Introduction

Proteins are critical molecular machines that underlie most biological processes and serve as foundational components in drug development, enzyme engineering, and disease research^1-3^. Their diverse functions arise from intricate conformational changes that allow them to interact both internally and with other molecules^4^. Consequently, elucidating the principles of protein folding and the ensemble of conformations is a prerequisite for interpreting protein function. Although experimental methods can probe the conformational ensemble and probability distribution with remarkable fidelity, they remain difficult to scale up. Nuclear magnetic resonance (NMR) can measure structural dynamics across a broad range of timescales, but the complex encoding of NMR data makes it challenging to resolve specific structural informations from the measurements^5^. Cryo-electron microscopy can provide multiple conformational states and their probability distributions, but is costly in both time and monetary expenditure^6^.

Molecular dynamics simulation (MD) is the principal computational strategy for exploring the conformational landscape of proteins. By numerically integrating Newton’s equations of motion, MD yields time-resolved trajectories that trace the positions of every atom within a molecular system and reveal conformational transition pathways, with the ensemble of sampled states conforming to Boltzmann distributions. Molecular dynamics simulations are categorised by resolution into all-atom and coarse-grained simulations. All-atom simulations retain every atom’s degrees of freedom within the system, enabling precise characterisation of intricate intramolecular dynamics^7-9^. However, computational demands for all-atom simulations exhibit non-linear scaling with atomic count, making sufficient sampling of conformational ensembles challenging. Conversely, coarse-grained force fields such as MARTINI^10^, CALVADOS^11^, and Mpipi^12^ map multiple atoms into a single coarse-grained particle, significantly reducing computational cost and enabling the simulation of larger biomolecular systems. However, coarse-grained methods sacrifice microscopic details within molecules and cannot describe precise atomic interactions.

Recently, advances in generative deep learning have dramatically accelerated structure–function studies of proteins, reducing the required computational time to just a few GPU hours^14-16^. Approaches such as the Boltzmann generator have demonstrated the potential of generative systems^17^, yet scaling these methods to larger proteins remains challenging. Methods for generating conformational ensembles can sample equilibrium protein conformations from implicitly learned Boltzmann distributions^18-21^, while machine learning-based MD models offer the ability to generate dynamical trajectories with reduced computational complexity^22-25^. Despite these advances, current generative methods still fall short of molecular dynamics simulations in sampling Boltzmann distributions, and models trained on MD data have yet to faithfully reproduce the conformational transition trajectories described by MD. Moreover, the computational cost of training conformational ensemble models also presents a significant challenge^18,21^.

Here, we introduced DynaFold, a robust latent diffusion-based generative framework capable of sampling conformational trajectories with high efficiency. Requiring training on only a small number of trajectories, DynaFold’s latent denoising model can capture protein dynamic trajectories and describe the transition paths between key conformations in a compute-friendly model size. Meanwhile, after removing temporal processing modules used for trajectories, DynaFold can also serve as a high-precision model for sampling conformational ensembles. DynaFold outperforms existing methods in conformational ensemble prediction and comformational changes modelling.

## Results

### Overview

We developed DynaFold, a generative framework to predict protein dynamics. DynaFold applies latent diffusion method that has been proven effective in computer vision for balancing computational efficiency with high-fidelity detail26. The core of DynaFold’s architecture is a Variational Autoencoder (VAE). This VAE includes an encoder that maps complex protein conformations into a simplified latent space, and a decoder that reconstructs conformations by sampling from this latent space. To model the latent diffusion process, we designed a Latent Denoising Transformer (LDT) as illustrated in Figure 1A. Trained on massive protein structures, DynaFold’s VAE organizes the conformational landscape into a more contiguous feature space. Within this space, LDT avoids the need of intricate model architectures to process complex three-dimensional structural representations^14,15^. Instead, it leverages only standard self-attention mechanisms to capture structural features and temporal dependencies in trajectories, thereby significantly enhancing computational efficiency.

**Figure 1.**
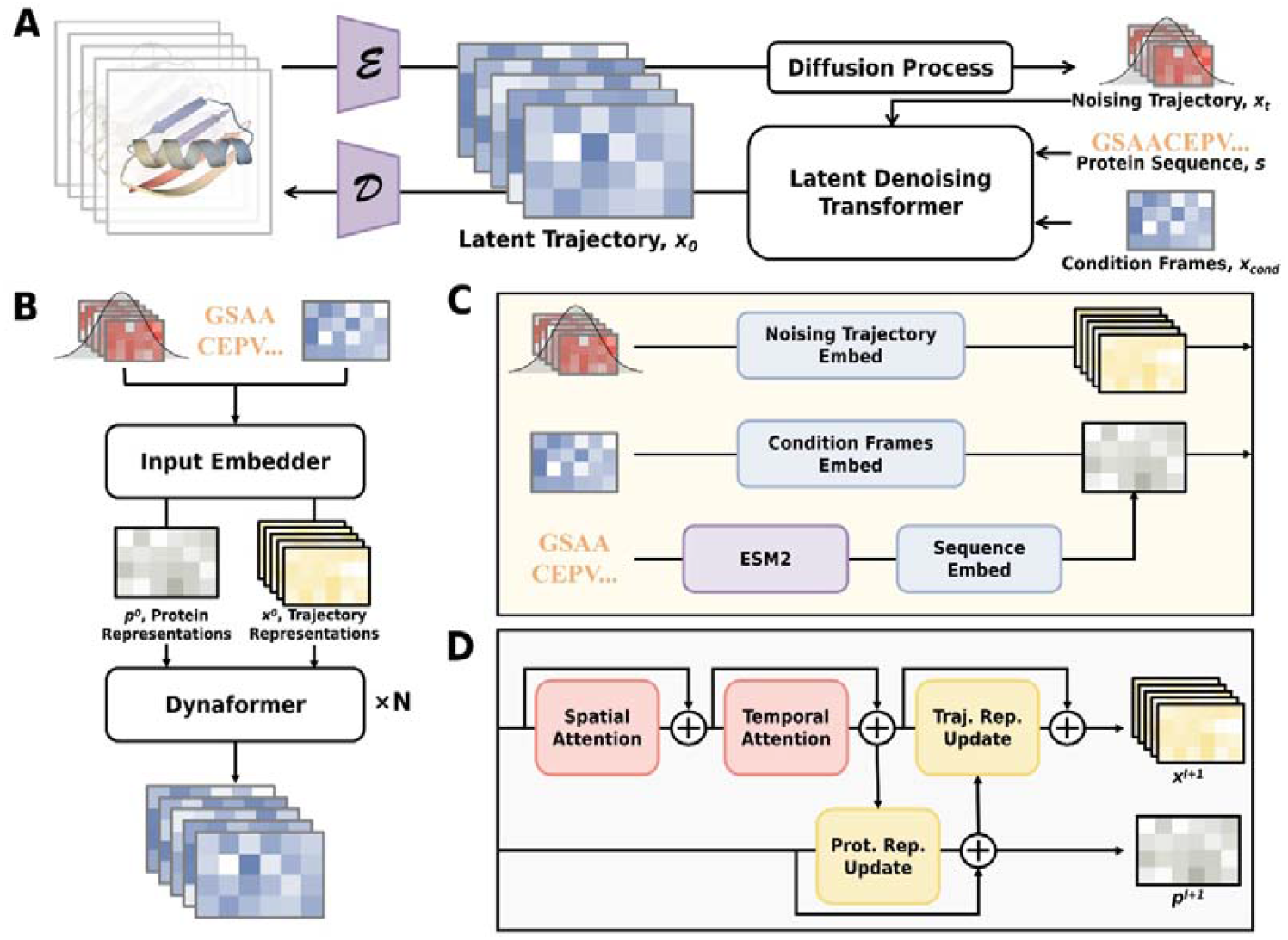
Overview of DynaFold. (A) Schematic representation of the DynaFold framework. E: backbone encoder. D: all-atom decoder. (B) Architecture of Latent Denoising Transformer. (C) Implementation details of Input Embedder. (D) Implementation details of Dynafomer.

DynaFold’s VAE adopts the model architecture introduced in ESM3^16^. Encoder accepts backbone structures and compresses each amino acid into an learned 12-dimensional feature representation. Decoder reconstructs these latent features into all-atom structures. LDT comprises an Input Embedder module and multiple Dynaformer modules as core processing components (Figure 1B). Input Embedder accepts noisy trajectories, conditioned on amino acid sequences and conditional frames, and processes all inputs into protein representations and trajectory representations (Figure 1C). Dynaformer iteratively updates both representations: trajectory representation first undergoes a spatial attention operation and a temporal attention operation to capture interactions in spatial and temporal dimensions, respectively. Subsequently, protein representation is updated, followed by trajectory representation (Figure 1D). Final layer of the Dynaformer’s trajectory representation directly reconstructs denoised trajectories.

We adopt EDM as the diffusion framework for DynaFold^27^. The training set consists of two parts: protein structures for VAE and molecular dynamics all-atom simulation trajectories for LDT. The structural data includes the Protein Data Bank (PDB)^28^ and AlphaFold Protein Structure Database (AFDB)^29^. The trajectory data includes ATLAS dataset^30^ and Fast-folding dataset^13^. DynaFold training is separated into two stages: in the first stage, a backbone resolution encoder-decoder is trained. In the second stage, the weights of backbone encoder are frozen, a all-atom decoder and LDTs are trained using the mean and variance of the latent space mapped by backbone encoder. Implementation details and training procedures are provided in Supplementary methods.

### DynaFold captures sub-microsecond-level dynamics

The Dynafold LDT model was initially trained on the ATLAS dataset to form a forward simulation model, wherein forward simulation refers to accepting an initial structure and predicting trajectories. We systematically evaluated DynaFold using 69 all-atom MD trajectories from ATLAS dataset^30^ that had never appeared during training. To fully capture the diversity learned by the diffusion model, we sampled three trajectories with 200 frames for each protein. We compare DynaFold with the all-atom state-of-the-art trajectory generation methods MDGen^22^, which have the same training set. MDGen accepts an initial structure and predicts dynamics trajectories, which has a similar prediction pattern and almost identical model scale to DynaFold (DynaFold’s LDT: 26M parameters, MDGen: 34M parameters).

We firstly calculated the average RMSF and pairwise RMSD of each case (Figure 2A, 2B). Dynafold captured,more comparable magnitude of conformational changes to MD simulations, indicating that our model can more accurately learn trajectory sequential features from training data. Meanwhile, DynaFold predicted noticeably larger RMSF and RMSD than MDGen predictions, demonstrating its capacity to sample more diverse conformations compared to models with similar scale.

**Figure 2.**
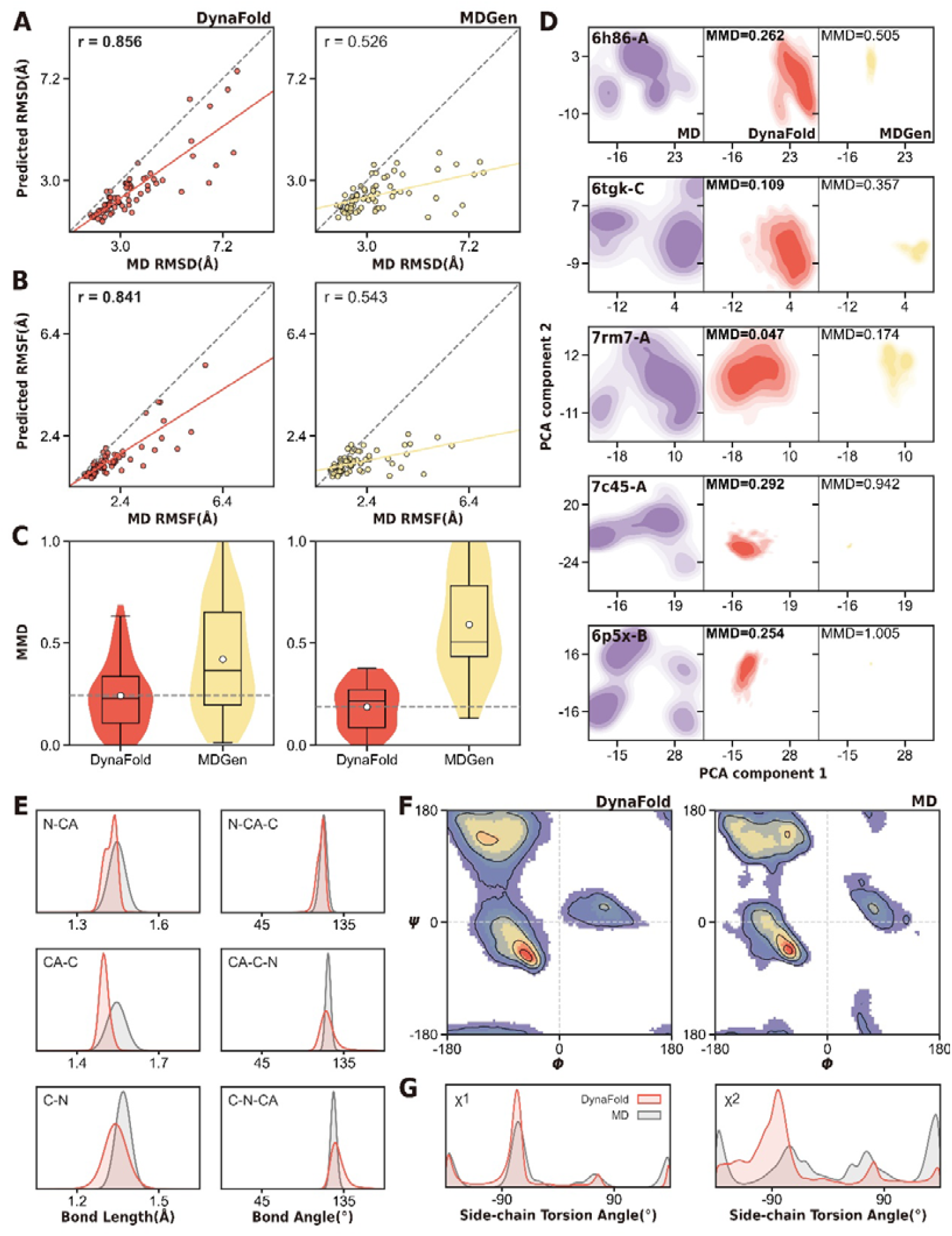
Performance of DynaFold for dynamics character. (A) All-atom pairwise RMSD between MD and model prediction, where pairwise RMSD denotes the average RMSD between any two frames within a trajectory. (B) All-atom RMSF between MD and model prediction. (C) Maximum Mean Discrepancies between MD and model prediction. (D) Distribution of protein trajectories with the maximum RMSF within each length interval predicted by each method in MD-fitted PCA space. (E) Backbone bond lengths and bond angles of MD and DynaFold prediction. (F) Backbone dihedral angles of MD and DynaFold prediction. (G) Side chain dihedral angles 𝒳_1_ and 𝒳_2_ of MD and DynaFold prediction.

**Figure 3.**
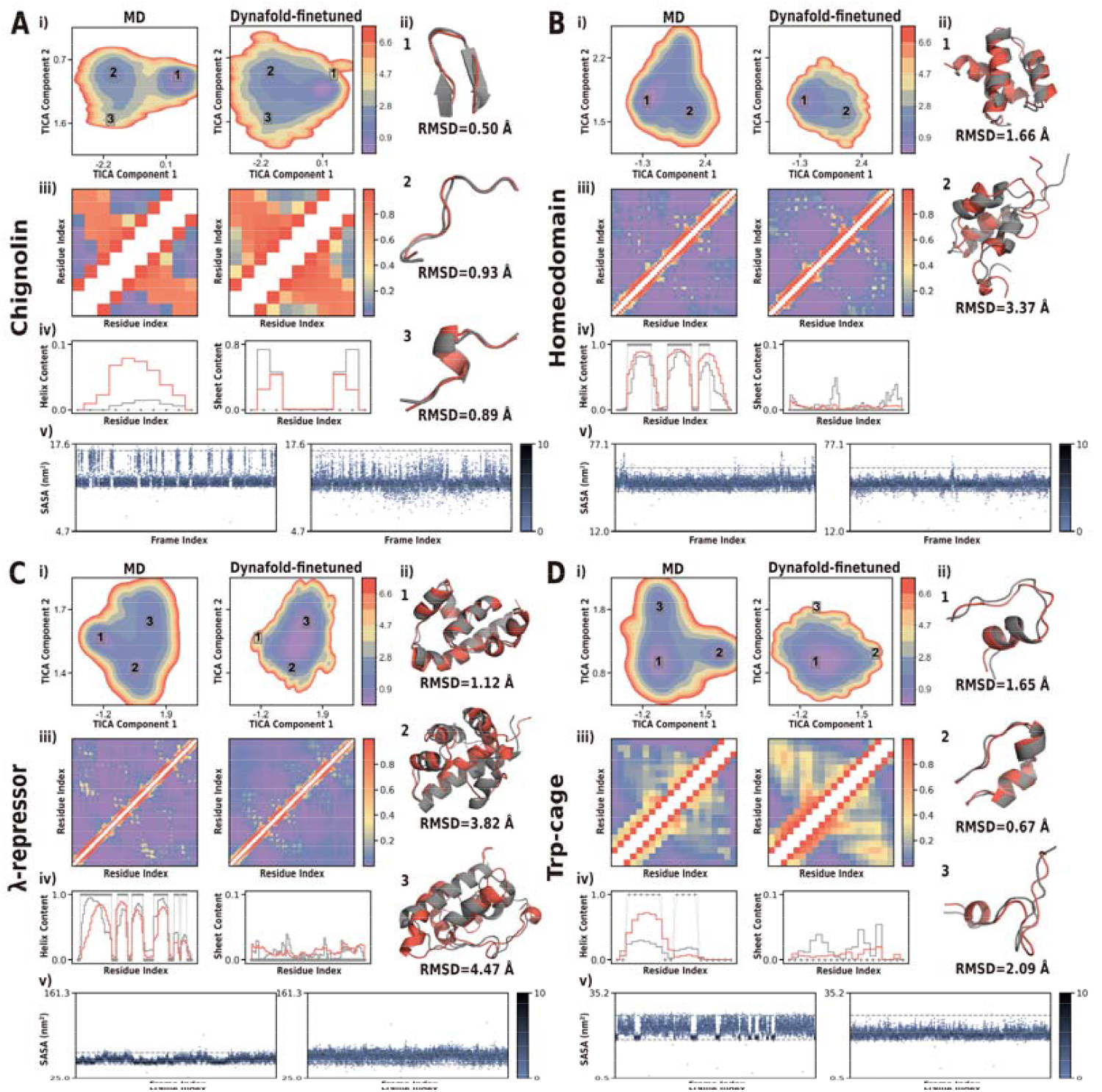
Performance of fine-tuned DynaFold on fast-folding proteins. Each subgraph consists of four parts to systematically evaluate the conformational ensemble and changes learned by DynaFold: i) MD ground truth and our model’s predicted free energy surface in MD-fitted TICA space. ii) Up to three local energy minimum conformations in the MD free energy surface and the model’s predicted minimum RMSD with them. Red: DynaFold. Grey: MD. iii) Residue contact frequency, defined as two residues with a Cα distance less than 8 Å. iv) Secondary structure frequency. Red: DynaFold. Grey solid line: MD. Grey dotted line: Initial experimental structures. v) The trajectory of solvent-accessible surface area (Top: MD, Bottom: DynaFold-finetuned). (A) Chignolin. (B) Homedomain. (C) λ-repressor. (D) Trp-cage.

Subsequently, we evaluated the consistency between model distributions and MD distributions in the test set in PCA spaces. We calculated the Maximum Mean Discrepancy (MMD)^40^ to quantify the distance between two distributions. DynaFold sampled ensembles that were more consistent to MD (Figure 2C Left). Since simulation time of trajectories in ATLAS dataset is relatively short, and conformational changes in most systems are modest, we further divided the test set into five intervals based on protein length and extracted the top three MD trajectories with the highest RMSF values from each interval (Table S1). In these 15 cases exhibiting greater conformational changes, DynaFold still outperformed MDGen in distribution consistency (Figure 2C Right, Figure S1, Table S2). In Figure 2D, we present cases with the largest RMSF in each interval. DynaFold captured a broader range of conformations across all cases, whereas MDGen tended to sample trajectories within a narrower conformational space or conservatively predict nearly identical conformations (shown as a single point in the PCA diagram). These results demonstrate that DynaFold’s architecture can capture molecular dynamics patterns more accurately.

Finally, We calculated the local geometric distribution of the conformations to examine the structural validity of trajectories predicted by DynaFold, including the bond lengths, bond angles, backbone dihedrals and side-chain dihedrals. DynaFold precisely reconstructed the typical backbone bond lengths and bond angle distributions, which closely matched the characteristics observed in the MD ensembles (Figure 2D). Additionally, Backbone dihedrals predicted by Dynafold fall within the acceptable region (Figure 2E). Moreover, DynaFold was also highly accurate in predicting χ1 (Figure 2F). All these metrics demonstrate that Dynafold can generate reasonable conformations. However, for amino acids with long side chains, the model could only capture some of the characteristics of χ2 and failed to predict the acute angle folding of χ3 and χ4 (Figure 2F, S2). Since the χ2-χ4 torsion angles are not uniquely determined by the backbone conformation and exhibit high diversity in the data distribution^43^, during training, the model is likely to prioritize fitting local means (χ2, χ3) or more frequently occurring features (χ4) to achieve the fastest decrease in loss, thereby getting stuck in a local minimum.

### DynaFold accurately estimates large-scale dynamics

The most robust and significant function of MD is its capability to exploring conformational ensemble of a system and sampling conformation change process. However, due to the simulation timescale limitations of ATLAS dataset, there is hardly significant conformational change in its trajectories. To further assess DynaFold’s ability to predict large-timescale simulations, we fine-tuned the forward simulation LDT using Fast-folding dataset from D. E. Shaw Research (DESRES) simulations^13^. Due to restrictions on DynaFold’s input, we removed one protein containing non-standard amino acids and trained the model with the remaining 11 proteins. Given the limited dataset size, we employed Leave One Out Cross Validation (LOOCV) for assessment, where a total of 11 models were used for model finetuning, with one protein retained for testing in each time and the remaining samples used for fine-tuning. Limited by model size, LDT struggles to completely fit complicated dynamic functions. Therefore, we uniformly sampled 10,000 frames from each trajectory to enable the model to learn rough dynamic trends over broad timescales.

For each protein, we first constructed a TICA^31,32^ space from MD trajectories to assess conformational ensembles predicted by our model. DynaFold predicted conformation ensembles comparable to those obtained by MD with only fine-tuning on 10 proteins (Figure 4i, Figure S3). However, constrained by model size and training data amount, free energy surfaces predicted by DynaFold is not accurate. For structural ensemble prediction methods, a key requirement is the capability to sample various stable states. To this end, we further assessed the model’s capability to capture key states by calculating the backbone RMSD between MD stable conformations and the most similar conformations predicted by the model. MD stable conformations here are defined as conformational points closest to low-energy points on the TICA diagram. DynaFold can sample various stable conformations in folded or partially folded states (Average RMSD = 2.68 Å) (Figure 4ii, Table S3). Moreover, as a crucial prerequisite for the realisation of protein function, secondary structures of non-loop regions predicted by Dynafold also exhibits comparable geometric consistency to that obtained via molecular dynamics simulations. (Figure 4ii, Figure S4, Table S4).

**Figure 4.**
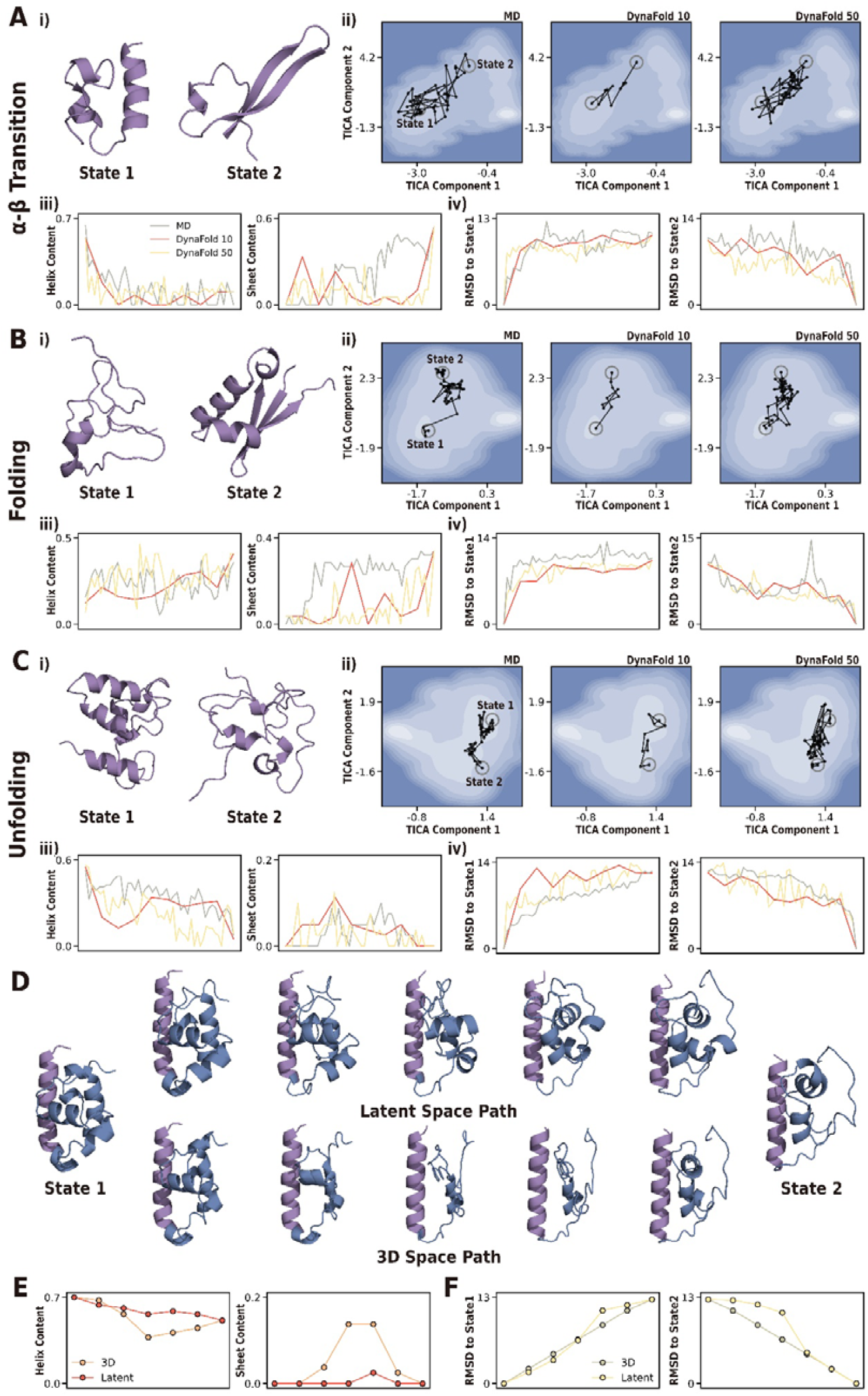
(A-C) Performance of different conformational transition events predicted by DynaFold on fast-folding proteins. The number following DynaFold denotes the frame count of sampling trajectories. Blue to white indicates a decrease in free energy. i) The initial (State 1) and final (State 2) structures of the conformational transition pathway. ii) MD ground truth and our model’s predicted pathways in MD-fitted TICA space. iii) Variations of secondary structures in pathways. iv) Backbone RMSDs between structures in pathways and States 1/2. (A) NLT9 transitions from an α-helix state to a β-sheet state. (B) ProteinG folds into a β-sheet dominated state. (C) λ-repressor unfolds from an α-helix state. (D-F) Comparison of continuity between our latent space and three dimensional space. (D) Interpolation path between two low-free-energy states of λ-repressor. (E) Secondary structure shifts in interpolated conformations. (F) RMSD between interpolated conformations and two states.

Subsequently, we examined the secondary structure distribution and residue contact frequency predicted by DynaFold. DynaFold effectively captures helix folding and overall residue contact patterns, but performs poorly on sheets (Figure 4iii,iv, Figure S5, Figure S6). In specific structures, we found that DynaFold can predict folds that are similar to sheets but cannot be definitively classified as sheets (Figure S4), indicating that the model still has limitations in sheet folding, which may be related to the extreme imbalance in secondary structure content in ATLAS and Fast Folding datasets. It is worth noting that DynaFold predicted more frequent contacts. Since the base model of DynaFold’s LDT is trained using trajectories composed mainly of highly structured conformations, which may have led to lower free energies for structured proteins in the conformational landscape modeled by DynaFold, causing the model that tends to sample compact conformations.

A major advantage of DynaFold over current protein ensemble prediction models is its ability to sample trajectory variation information. Therefore, we calculated the solvent accessible surface area (SASA) of MD and DynaFold-predicted trajectories to assess DynaFold’s capacity for capturing temporal sequence information of structural folding. The SASA predicted by DynaFold falls within an acceptable range, indicating that the model effectively captures main features of protein folding and unfolding processes. Notably, compared to conventional MD simulations, DynaFold exhibits more frequent conformational transitions while reducing repetitive motion within stable conformational states. Although this limits DynaFold’s accuracy in free energy prediction, it enables more efficient exploration of protein structural ensembles (Figure 4vi, Figure S7).

### DynaFold precisely samples continuous conformational transition pathways

Exploring conformational transition pathways between two specific conformations holds great significance for investigating the functional formation and transition of proteins. However, this remains challenging for MD methods and ensemble prediction models. To this end, we further fine-tuned DynaFold’s LDT using the Fast-folding dataset to enable sampling of transition trajectories between two conformations at different time steps (see Materials and Methods). During training and assessing, each trajectory was uniformly downsampled to 100,000 frames. We extracted three distinct events from test protein trajectories to evaluate DynaFold’s performance in sampling different types of conformational changes: secondary structure type transitions (α-β transition), folding and unfolding. For each event, we sampled 10-frame and 50-frame trajectories to evaluate DynaFold’s capability in predicting conformational transition trajectories across different time scales.

We compared conformational transition pathways predicted by DynaFold and MD simulations in the MD-fitted TICA space, alongside conformational change characteristics in three-dimensional space. The three-dimensional conformational characteristics include secondary structure changes and backbone RMSDs between model predicted tracjectories and their corresponding initial/final structures. DynaFold predicted TICA pathways landing in areas close to MD simulations (Figure 4ii), with secondary structure changes occurring within acceptable ranges (Figure 4iii). Meanwhile, RMSD variation trends in model predicted trajectories are consistent with MD simulations (Figure 4iv), indicating that DynaFold is capable of capturing temporal dynamics of trajectories and predicting continuous transition pathways between two distinct conformations.

Specifically, for the α-β transition event, both MD and DynaFold initially sampled conformations within the low-free-surface region around State 1, and approached State 2 with comparable RMSD trends. Both methods predicted the unfolding of α-helix at the beginning of trajectories, but the formation of complete β-sheet occurred later than in MD simulations (Figure 4A). For the folding event, both MD and DynaFold predicted conformations moving away from State 1 at the start of trajectories in TICA space, with most conformational points aggregating closer to State 2. DynaFold trajectories maintained α-helix levels consistent with MD during β-sheet folding and predicted β-sheet formation similar to MD early in transition pathways. However, the formed β-sheet could not be stably maintained like in MD, instead reverting to an unfolded state before refolding (Figure 4B). For Unfolding events, both MD and DynaFold avoided sampling conformations in high-free-energy regions along transition pathways. Although DynaFold predicted greater fluctuations in α-helix content during unfolding, the overall trend aligned with a gradual unfolding process, and the variation in β-sheet content also fell into acceptable ranges (Figure 4C).

It is worth noting that, as TICA dimension reduction places more emphasis on mapping dynamically fast-equilibrium conformations (those frequently interconverting during simulations) to neighbouring positions in the TICA space^32^, whereas the structural points of conformational transition trajectories predicted by DynaFold consistently lie in proximity to MD trajectories, with spans between any two points remaining within comparable ranges. This demonstrates DynaFold’s capability to capture the conformational change characteristics described by dynamics at each time step. Moreover, owing to the stochastic nature of MD, transition pathways between any two conformations are not unique. DynaFold is capable of capturing diverse transition pathways between two states, and sampled conformations tend to aggregate near low free energy regions (Figure S8). This indicates that DynaFold can also explore conformation ensembles and estimate probability distributions along these transition pathways.

During evaluation process, we found that DynaFold generally captures more accurate distributions than comparable models, and we argue that this is related to the spatial discretization of the data. The three-dimensional structure of proteins is not continuously distributed in three-dimensional space but exhibits considerable discretization. This discretization stems from the strict physical-chemical constraints between atoms, rendering most points in three-dimensional space physically unfeasible. Such a discrete data distribution makes it more challenging for models to capture structural correlations. In contrast, our VAE with KL divergence loss during training maps proteins into a more continuous space. To demonstrate the impact of DynaFold’s VAE on data distribution continuity, we selected two low-free-energy conformations of λ-repressor. After aligning 20 highly similar helix residues at the N-terminus (purple region), we performed interpolation between the two conformations in both the latent space and the three-dimensional space, and assessed conformational features along the interpolated transition path. For the three-dimensional space, we performed linear interpolation directly. For the latent space, since the VAE was trained with a large variance, regions near the VAE-predicted mean were still decoded as highly similar structures. Therefore, for the interpolated conformation x_i_ = 1 − w_i_ · state_1_ + w_i_ · state_2_, we set the weights w_i_ ∈ [0.35,0.45,0.5,0.55,0.65].

We first presented the conformational transition pathways. The conformations interpolated in the latent space all maintained reasonable folding, whereas in 3D space, folding that clearly violated physical constraints was observed (Figure 5D). We subsequently calculated the secondary structure content of the interpolated conformations and the RMSD between the two states (Figure 5E, 5F). The latent space interpolated conformations exhibit a more linear change in secondary structure content, while interpolation in three-dimensional structures results in a sudden increase in sheet content that is incongruous with helix-dominated conformations. The change in RMSD also indicates a nonlinear relationship between latent space and three-dimensional space. This demonstrates that our VAE indeed learned the functional similarity described by three-dimensional structures and mapped chemically meaningful and related protein structures to more adjacent positions, thereby improving the continuity of the data distribution.

**Figure 5.**
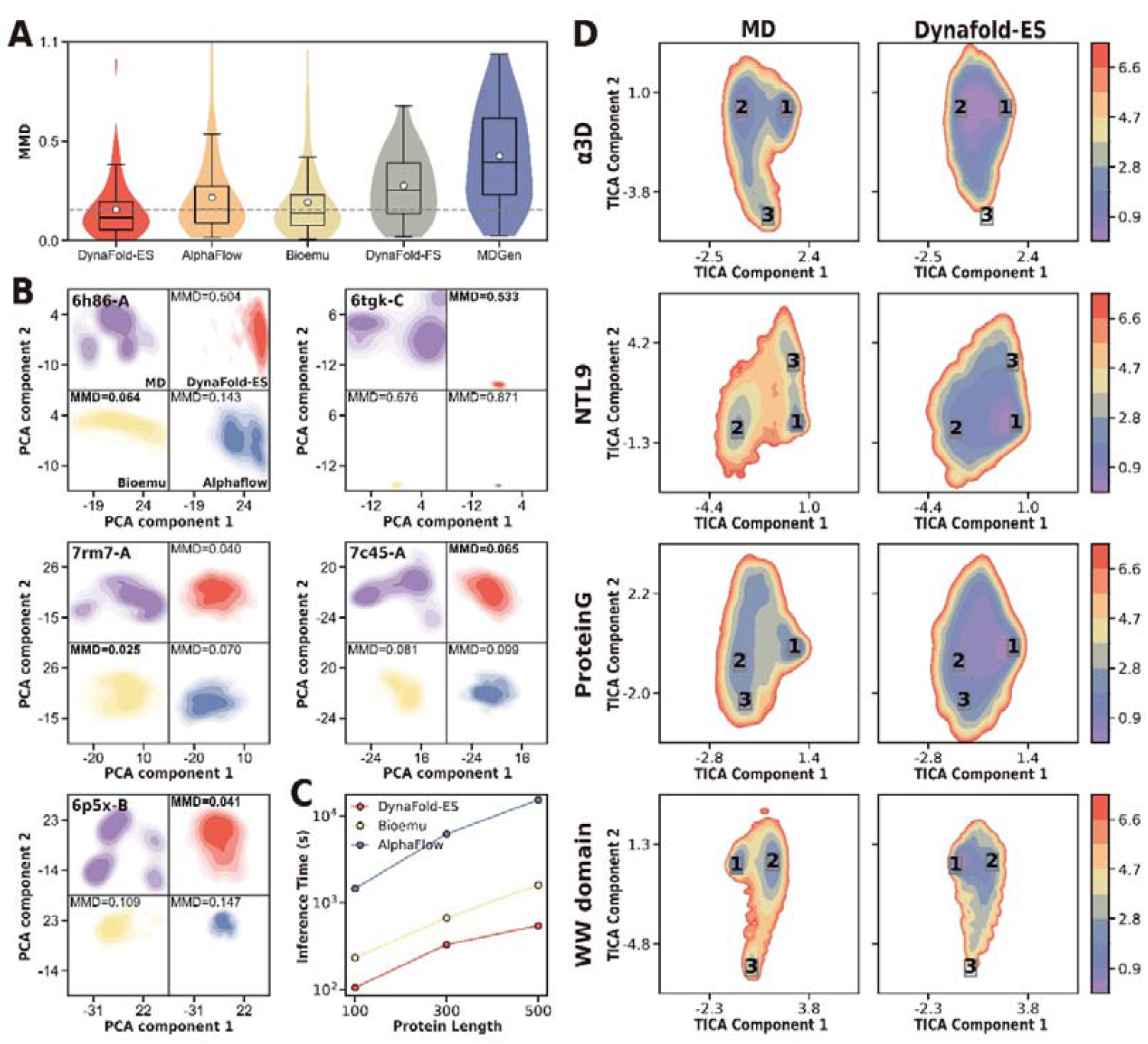
Performance of DynaFold-ES on different MD datasets. (A) Maximum mean discrepancies between model predictions and MD on 69 ATLAS test proteins. White dots denote the mean values. (B) Distributions of protein ensembles with the maximum RMSF within each length interval predicted by each method in MD-fitted PCA space. (C) Computational time of sampling 200 structures for different protein lengths. (D) Free energy surface of MD ground truth and our model’s prediction in MD-fitted TICA space.

### DynaFold efficiently generates structural ensembles

Forward simulation DynaFold is constrained by its limited computational complexity, making it difficult to capture the complete dynamic information within training set. To further evaluate DynaFold’s capability in sampling conformational distributions, we removed temporal attention modules and conditional structures input from LDT and expanded the model size to train a sequence-to-ensemble model. As ensemble models require learning complex sequence-to-structure mapping functions, we pre-trained DynaFold using the VAE training dataset. Prior to training, we applied a two-step clustering process to balance sequence distributions: 1) 90% similarity clustering, retaining only cluster centres. 2) 40% similarity clustering, with one sample randomly selected from each cluster per epoch during training. During the evaluation process, we designated the ensemble prediction model and the forward simulation model as DynaFold-ES and DynaFold-FS respectively.

We firstly fine-tuned DynaFold-ES’s LDT on the same ALTAS training set^21^ and compared it against state-of-the-art methods AlphaFlow^21^ and Bioemu^18^ on the test set. During evaluation, each model sampled 600 structures per protein. We evaluated conformational distributions in MD-fitted PCA space. As Bioemu serves as the backbone model, we employed only atoms predicted by Bioemu for dimensionality reduction during PCA. DynaFold-ES predicted conformational distributions most closely resembling MD (Figure 5A), and among the 5 cases exhibiting the largest RMSF in each interval, DynaFold demonstrated closest distributions in 3 cases (Figure 5B). Concurrently, DynaFold demonstrated the highest computational efficiency among all methods, achieving a tenfold reduction in computation time compared to AlphaFlow, another all-atom approach (Figure 5C). These results indicate that DynaFold outperforms existing methods in both accuracy and efficiency of capturing structural ensembles described by molecular dynamics.

Subsequently, we fine-tuned the pre-trained DynaFold-ES’s LDT on Fast Folding dataset. During both training and assessment, each trajectory was uniformly sampled to 100,000 frames to capture more detailed information about dynamical distributions. Evaluation was conducted on four proteins where forward simulation LDTs performed poorly. DynaFold-ES captured a consistent range of conformations and successfully predicted the low free energy of stable conformations (Figure 5D, Table S5), indicating that DynaFold, after scaling up and pre-training on structural data, can capture dynamics features from trajectories of just ten proteins. However, DynaFold tends to underestimate the free energy in regions between stable conformational points, indicating that relying solely on implicit modelling of conformational probabilities presents significant challenges. Achieving high-precision predictions of conformational free energy requires either further expanding the model size or incorporating explicit objective functions to guide the model. Furthermore, we fine-tuned DynaFold using the same 10,000-frame data as forward simulation models. DynaFold-ES exhibited a free energy surface more consistent with MD simulations compared to DynaFold-FS (Figure S9, Table S6), demonstrating that DynaFold-ES’s performance improvement stems from the combined effect of model and data scaling up.

## Discussion

We developed DynaFold, a latent diffusion based generative framework for predicting all-atom trajectories and conformational ensembles of proteins. This framework has demonstrated its strong performance in sampling protein conformational ensembles and transition pathways.

The major difference between DynaFold and other ensemble prediction methods is its architectural innovation. The success of traditional structural or ensemble prediction models strongly relies on mapping multiple sequence alignment (MSA) information to three-dimensional structural space through complex model structure. In contrast, DynaFold reduces the complexity of the data space through a Variational Autoencoder (VAE) and accepts initial backbone structures or single sequence preprocessed by protein language models as inputs to eliminate dependence on MSA, thereby lowering the difficulty of model training. Ultimately, DynaFold outperforms existing methods in both trajectory and ensemble prediction with lower computational complexity. Meanwhile, DynaFold is capable of capturing conformational transition pathways that ensemble prediction models cannot describe, providing insights into the unfolding and folding processes between stable conformations.

Training protein structure models is often computationally intensive. Many methods have been proposed to convert protein structures into different representations to simplify the complexity of modelling protein energy landscapes^16,33^. We introduce latent diffusion models into conformational trajectories generation, essentially hoping to provide further insights into lightweight learning. However, the success of this method highly depends on reducing the complexity of the latent space while maintaining high precision in structure reconstruction. Currently, DynaFold’s VAE does not perform perfectly in structure reconstruction, failing to accurately reconstruct a small portion of sheet structures and some atoms in long side chains. Additionally, our VAE was trained on experimental crystal structures, and predictions cannot capture accurate side-chain dynamics. Therefore, for studies requiring precise side-chain conformations, all-atom simulations remain the more recommended choice.

DynaFold has certain limitations in accurately predicting free energy surfaces. Constrained by the limited number of long-simulation all-atom trajectories available for training, even state-of-the-art models cannot accurately predict free energy surfaces of proteins^18^. For the rapid exploration of approximate conformational free energy surfaces, coarse-grained force fields remain the optimal choice. For deep learning approaches aiming to achieve high-precision modelling of free energy surfaces for full-atom conformations, further development may be required in the following two areas: 1) Designing efficient and accurate free energy objective functions. 2) Pre-training models on extensive coarse-grained force field trajectories and crystal structures, then transferring them to all-atom resolution.

## Materials and Methods

### Dataset

The training set for DynaFold’s VAE consists of high-quality crystal structures from the Protein Data Bank (PDB)^28^ and predicted structures from the AlphaFold Protein Structure Database (AFDB)^29^. While the training set for LDT consists of two all-atom simulation datasets, ATLAS^30^ and Fast Folding^13^.

For crystal structures, we gathered all X-ray-resolved protein chains in PDB released before 2025-01-01, applying filters: (1) resolution ≤ 8Å, (2) minimum of 5 residues, (3) retaining only one chain per entity ID. Structures released before 2023-09-30 form training set, while those released after 2023-10-01 constitute test set. To rigorously assess model generalization, we used mmseqs2^34^ for clustering (parameters: --min-seq-id 0.4 --cov-mode 1 --cluster-mode 2 --kmer-per-seq 100 -c 0.8), retaining only test set proteins with less than 40% sequence similarity to training set. Additionally, during testing, we exclusively employed proteins with lengths ranging from 30 to 1000 amino acids. This yield 231,751 training structures and 385 test structures.

For AlphaFold predicted structures, we utilized a subset from AFDB v4, encompassing 48 species proteomes and most Swiss-Prot proteins. To ensure training data quality, we selected only predicted structures with global pLDDT exceeding 0.7. Additionally, to address predominance of alpha-helix structures in AlphaFold predictions and balance secondary structure distribution, we measure long-range contacts within protein chains. A long-range contact is defined as Cα atoms of two amino acids, separated by more than 12 residues, with physical distance less than 8 Å. Proteins of length L with fewer than 0.5L long-range contacts are excluded^16^. This process yield an additional 704,391 training structures. During training process, We randomly select 2,000 structures each from PDB and AFDB datasets to constitute validation set.

For ATLAS dataset, we adopt same training and test set partitioning as AlphaFlow^21^. For the test set, only proteins shorter than 500 are retained, finally comprising 1,267 trajectories for training, 39 for validation, and 69 for testing. To enable models to explore a broad conformational space more efficiently, we sample one frame every 500 ps, resulting in each trajectory consisting of 201 frames.

### Latent Diffusion

We employed EDM^27^ as the diffusion framework to learn mapping from Gaussian noise distribution to protein conformation equilibrium distribution in the latent space. Since each protein in the latent space is represented as two-dimensional matrix [protein_length, feature_dimension], we do not need to adapt diffusion algorithm for protein structural data, unlike previous protein structure diffusion models. The distribution of noise samples at the forward

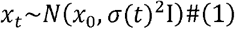

diffusion process can be formulated as

Where *σ*(*t*) denotes a noise schedule function used to control the noise intensity at each diffusion timestep.

The reverse denoising process can be formulated as an ordinary differential equation (ODE). Specifically, the EDM method introduces a generalized ODE representation, which can be expressed as follows

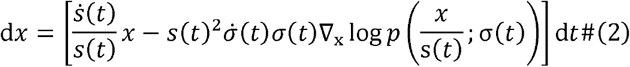

Where *s*(*t*) denotes a scale schedule function used to control the scaling factor applied to the data at each diffusion time step. ∇_x_ log *p* denotes the score function, defined as the gradient of the log-probability density function with respect to input data.

The EDM method trained a network *F*_*θ*_(*x,t*) and utilized *F*_*θ*_(*x,t*) to derive the denoiser D. However, we found this approach ineffective for latent protein trajectory reconstruction during training. Consequently, we directly parameterized denoising network *D*_*θ*_(*x,t*) to learn the reverse diffusion process. Since the diffusion time step *t* and the noise level *σ* are interconvertible, the EDM framework samples σ directly from a given noise distribution during training instead of sampling a *t* and calculating *σ*. General form of the objective function for *D*_*θ*_(*x,t*) is expressed as follows (see Appendix 4.2.1 for specific loss function description)

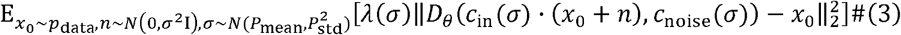

Where *n* is noise, *p*_data_ is data distribution, *P*_mean_ = − 1.2 and *P*_std_=1.2 determine the noise distribution during training. *c*_in_ scales inputs with different noise levels, *c*_noise_ maps noise levels to conditional inputs of *D*_*θ*_, and *λ*(*σ*) weights expectations over different noise levels. These three parameters adopt identical settings as in EDM^27^.

### VAE Architecture

DynaFold’s VAE encoder adopts the architecture from VQ-VAE of ESM3^16^ and replaces the discrete codebook with continuous mean and variance outputs, enabling compatibility with diffusion processes in continuous latent space. The decoder consists of stacked Transformer layers employing standard self-attention operations^36^, and is designed to directly predict the Cartesian coordinates of amino acid atoms.

### VAE Loss Function

The details of the loss function for VAE training are as follows, consisting of an all-atom mean square error (MSE), a smooth local distance difference test (lDDT)^37^, a side-chain pairwise distance loss, a bond length penalty and a Kullback-Leibler (KL) divergence.

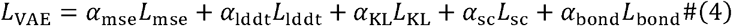

Where *α*_mse_ = *α*_lddt_ = 1, *α*_bond_ = *α*_KL_ = 0.2. At the start of training, *α*_sc_ = 0; when the loss approaches convergence, *α*_sc_= 1.

After performing a rigid alignment between the predicted structure 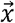 and the ground truth 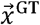, we calculated all-atom MSE loss as

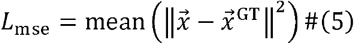

We employed the smooth lddt as an auxiliary structure reconstruction loss as

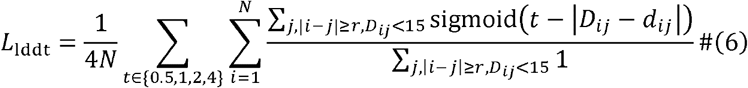

Where *N* denotes the number of protein residues, *D*_*ij*_ denotes the distance between Cα atoms within 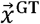 and *d*_*ij*_ denotes the distance between these pairs of atoms within 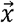.

In order to reduce the occurrence of clash structures, we calculated a bond length penalty as

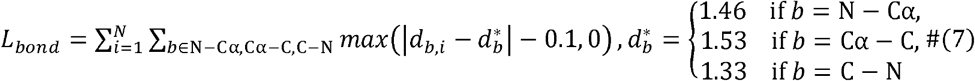

Where *N* denotes the number of protein residues, *d*_*b,i*_ denotes *b* bond of the i-th residue.

Since protein structure models tend to prioritize learning backbone conformations during training and stack side chain groups near backbone atoms to achieve maximum loss reduction efficiency, we added a side chain pairwise distance loss when the loss approached convergence to emphasize the significance of side chain conformations to the model. This loss term is calculated as

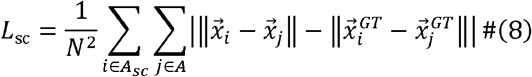

Where *A* and *A*_sc_ represent the all-atom set and side-chain atom set of a protein, 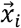 denotes the coordinate of *i* atom in the predicted structure, 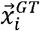 denotes the coordinate of *i* atom in the ground truth.

We calculated a KL divergence loss to ensure the model learns a more reasonable latent space distribution, and we define the target distribution as a

Gaussian distribution with μ_target_ = 0 and 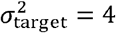. Choosing a larger variance than the standard Gaussian distribution is due to the fact that the denoising results of the diffusion model may be shifted from the true mean, and setting target variance to a larger value can improve the decoder’s prediction accuracy when facing shifted samples. We calculate the KL divergence loss as

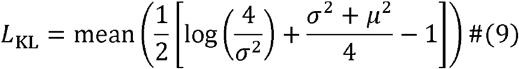

Where *μ* and *σ* _2_ denote the mean and variance of the encoder’s prediction.

### VAE Implementation Details

The training of VAE consists of two stages: backbone encoder-decoder (stage 1) and all-atom decoder (stage 2). In Table 1, we presented the model hyperparameters utilized during the two training stages. During both training stages, we used the AdamW optimizer^38^ with learning rate warming up over 5,000 steps to 5e-4. Once training loss ceases to decrease, we applied cosine decay to anneal learning rate to 5e-5. Maximum sequence length of proteins were cropped to 512. All stages of VAE training were conducted on four NVIDIA A100 40GB PCIe. Both stages were trained on 4 NVIDIA A100 40GB PCIe. In Table 2, we present the average all-atom RMSD and Cα TM-score between DynaFold’s VAE-reconstructed structures and the ground truth on the PDB test set.

**Table 1.** Hyperparameters of VAEs.

| Hyperparameters |  | Stage 1 | Stage 2 |
| --- | --- | --- | --- |
| Encoder | Geometric Attention Layers | 2 | 2 |
|  | Geometric Attention Heads | 128 | 128 |
|  | Hidden Dim | 1024 | 1024 |
|  | Latent Dim | 12 | 12 |
|  | Freeze Weights | False | True |
|  | Model Parameters | 31M | 31M |
| Decoder | Structural Resolution | Backbone | All Atom |
|  | Transformer Blocks | 8 | 12 |
|  | Attention Heads | 16 | 16 |
|  | Hidden Dim | 1024 | 1024 |
|  | Model Parameters | 106M | 158M |

**Table 2.** Metrics of VAE reconstructed structures.

| Length range | All-Atom RMSD (Å) | TM-Score |
| --- | --- | --- |
| (0,100] | 2.118 | 0.972 |
| (100,300] | 1.545 | 0.983 |
| (300,600] | 1.663 | 0.987 |
| (600,1000] | 1.510 | 0.993 |
| All | 1.735 | 0.981 |

### LDT Architecture

Here, for other variables that appear in the algorithm, we use the following notation:

- *T*: number of trajectory frames
- *L*: number of residues
- *t*: diffusion time step
- *m*: condition mask

In Supplementary algorithms 1–3, we present our novel model architecture, Latent Denoising Transformer (LDT), tailored for protein latent trajectory diffusion tasks. LDT integrates an Input Embedder module with multiple Dynaformer modules. Within LDT, latent noisy trajectory *x*_*t*_, conditional frames *x*_cond_ and amino acid sequences s are processed into protein representations *p*^*l*^ and trajectory representations *x*^*l*^ (Superscript *l* denotes output of layer l), both iteratively updated by Dynaformer. This innovative feature representations enables network to capture comprehensive protein description and detailed differ initiation of individual structures within a trajectory in a detailed manner. The final layer of Dynaformer’s trajectory representation is used to predict latent trajectories *x*_0_.

The process of generating protein representation and trajectory representation by Input Embedder from input data is as follows:

1. **Sequence Embed:** Amino acid sequences are initially processed by ESM2 650M^39^. Token representations *s*_*t*_ for each residue are derived by averaging representations across 34 layers. Sequence representations *s*_*s*_ are then obtained by averaging all token representations. Both representations, along with original amino acid sequences, are linearly projected and summed to form sequence component of protein representations *p*_*s*_.
2. **Condition Frames Embed:** Conditional frames *x*_cond_ are initialised with the same shape as the noise trajectory, where all values are 0 except for the position of the conditional frame (in Dynafold, only the first frame is a conditional frame). Subsequently, conditional frames and condition masks are linearly projected and summed to yield condition representations *c*. Meanwhile, conditional frames are also processed through another linear projection and averaging to derive conformational component of protein representations *p*_*c*_.
3. **Representation construction:** Protein representations is derived by summing its sequence component and conformational component. Noisy trajectories, following linear projection, is combined with protein representations and condition representations to yield trajectory representations.

Dynaformer updates the protein representations and trajectory representations as follows:

1. **Diffusion time step scaling:** In each Dynaformer module, diffusion timestep is linearly projected into scaling and bias parameters, which incorporate diffusion timescale information during attention and MLP operations of trajectory representations.
2. **Attention Operations:** Attention operations are sequentially applied to each conformational interior (spatial dimension) and each residue time series (temporal dimension) of trajectory representations. Both temporal and spatial attention module employ rotational position encoding^22^ (RoPE).
3. **Representations Update** : Protein representations and mean of trajectory representations are first concatenated and, after linear projection, added to protein representations. Protein representations are then updated by a MLP using gaussian error gated linear units (GEGLU)^40^ as the activation function. Trajectory representations are updated directly through a GEGLU-MLP after adding the updated protein representations.

### LDT Loss Function

The details of the loss function for LDT training are as follows, consisting of a mean square error (MSE) and a pairwise distance (PD) loss. Comparing to our denoising reconstruction objective function (Equation 2) given in Latent Diffusion, we additionally added the pairwise distance loss as an auxiliary loss to help the model capture more details during the training process.

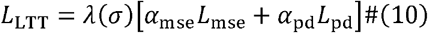

Where *α*_mse_ = 1, *α*_pd_ = 0.2.

We calculated the MSE loss as

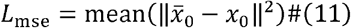

Where 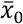 denote the predicted denoised sample of latent space trajectory and *x*_0_ denotes the ground truth.

We employed the pairwise distance loss as an auxiliary denoising reconstruction loss as

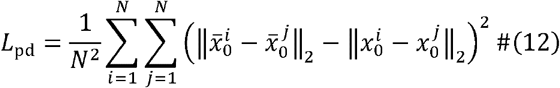

Where *N* denotes the number of protein residues. The superscript *i* or *j* denotes the latent features of the the i-th or j-th residue.

### LDT Implementation Details

Table 3 presents model hyperparameters utilized for both LDT training and fine-tuning. Table 4 presents training details utilized for different training stages, where the forward simulation, conformational transition and ensemble models are abbreviated as FS, CT and ES respectively. We employ AdamW^38^ as the optimizer. During model training using Fast Folding dataset, owing to the extreme imbalance in α-helix and β-sheet content within the dataset^13^, we added an additional loss weighting factor during training. This factor, numerically equal to β-sheet contents of structures, was applied to all frames with β-sheet content exceeding 0.2 in a trajectory. During training of the conformation transition model, to ensure that LDT can sample conformation transition pathways with with varying degrees of conformational change and different time step, we randomly sample sub-trajectories with *n* frames per epoch for each sample and then randomly downsampled these sub-trajectories to 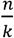 frames (*n* ∈ [50,500] ∩ ℤ and *k* ∈ [1,5] ∩ ℤ). All stages were trained on 1 NVIDIA A100 40GB PCIe.

**Table 3.** Model hyperparameters of LDT.

| Hyperparameters | FS/CT | ES |
| --- | --- | --- |
| Dynaformer Blocks | 6 | 12 |
| Attention Heads | 6 | 16 |
| Hidden Dim | 384 | 1024 |
| MLP Hidden Dim | 768 | 2048 |
| Use Condition Frames | True | False |
| Model Parameters | 26M | 306M |

**Table 4.**
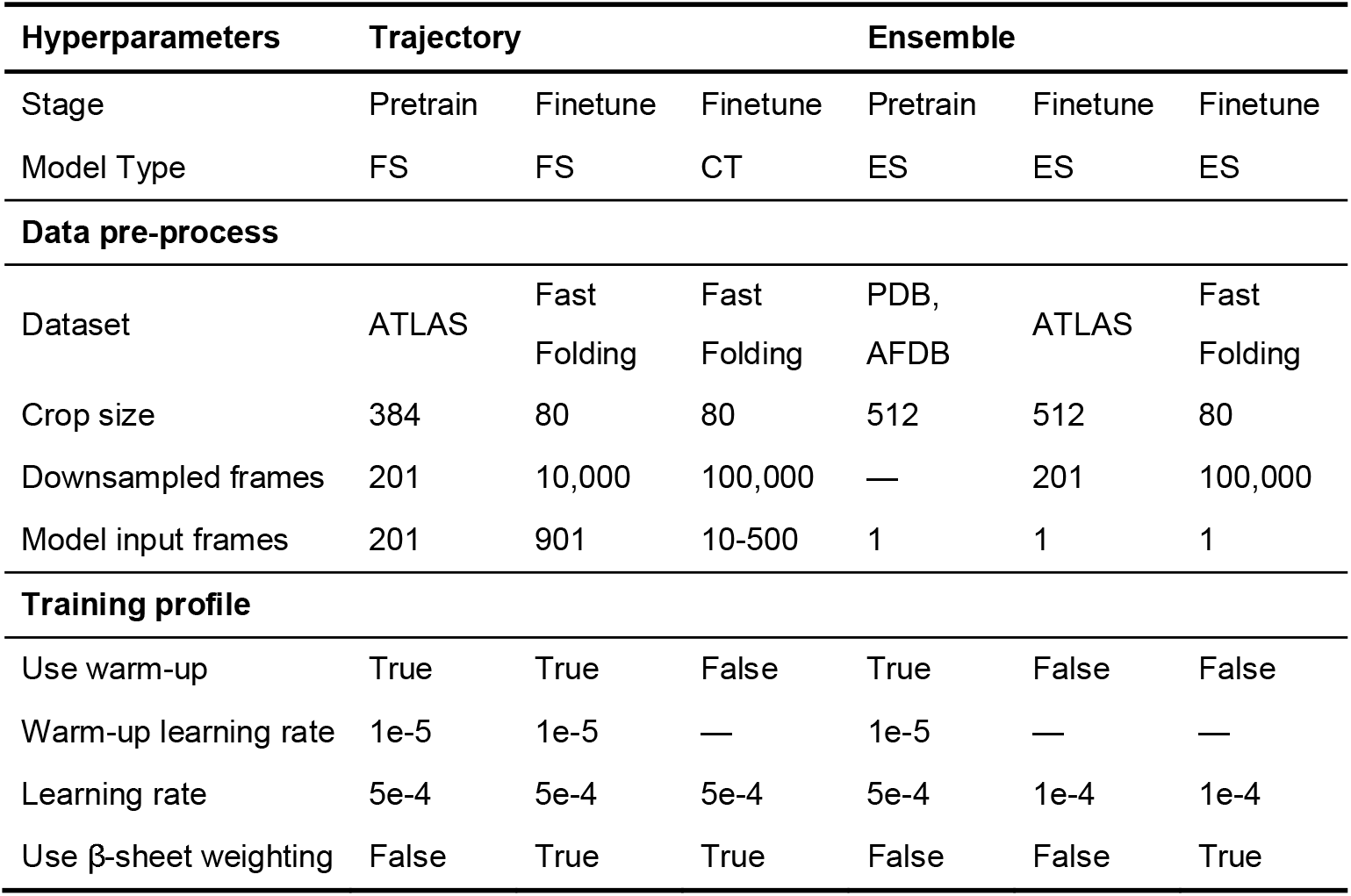
Training details of LDT.

| Hyperparameters | Trajectory |  |  | Ensemble |  |  |
| --- | --- | --- | --- | --- | --- | --- |
| Stage | Pretrain | Finetune | Finetune | Pretrain | Finetune | Finetune |
| Model Type | FS | FS | CT | ES | ES | ES |
| **Data pre-process** |  |  |  |  |  |  |
| Dataset | ATLAS | Fast Folding | Fast Folding | PDB, AFDB | ATLAS | Fast Folding |
| Crop size | 384 | 80 | 80 | 512 | 512 | 80 |
| Downsampled frames | 201 | 10,000 | 100,000 | — | 201 | 100,000 |
| Model input frames | 201 | 901 | 10-500 | 1 | 1 | 1 |
| **Training profile** |  |  |  |  |  |  |
| Use warm-up | True | True | False | True | False | False |
| Warm-up learning rate | 1e-5 | 1e-5 | — | 1e-5 | — | — |
| Learning rate | 5e-4 | 5e-4 | 5e-4 | 5e-4 | 1e-4 | 1e-4 |
| Use β-sheet weighting | False | True | True | False | False | True |

### ATLAS Evaluation Metrics

In ATLAS^30^ test set, we compare the similarity between generated ensembles and MD ensembles in two aspects: (1) quantitative assessment and (2) structural validity. For evaluation, MD trajectories were sampled at intervals of 500 ps to match the dynamic scale learned by our model at this time step during training. In each inference, the trajectory generation models accepted an initial structure and subsequently sampled the dynamic trajectories of 200 frames. The conformational ensemble generation models, by contrast, directly generated 200 independent structures. During the reverse diffusion process, we set the maximum and minimum noise levels to σ_max_ = 80 and σ_min_ = 0.001, respectively, and performed denoising across 500 steps; the reverse ODE was numerically integrated using the Heun second-order solver. All other baseline models were run with their default parameter settings^21,22^. Furthermore, for consistency and comparability with the 3 repeated MD trajectories available for each protein in the ATLAS dataset, we sampled each protein three times for each model.

For quantitative assessment, we initially calculated the average pairwise RMSD and RMSF for each sample in each method to preliminarily verify the conformational diversity captured by models and MD. Where both RMSD and RMSF are calculated on all atoms of proteins, and pairwise denotes calculations between any two frames within a trajectory. Subsequently, we divided the length of 500 into five equal intervals and extracted the three proteins with the highest RMSF in each interval. We then specifically evaluated the similarity between distributions of model-predicted trajectories and MD-predicted trajectories for each protein in Rg-RMSD space and PCA space. The similarity between the two distributions was examined using Maximum Mean Discrepancy (MMD)^40^, which can be calculated as

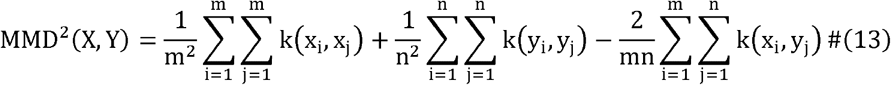

Where k(x,y) = exp (− γ ∥x − y∥^2^ is a kernel function used for feature mapping, 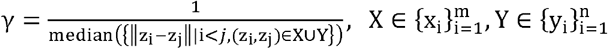 denotes the point set under the current space of two methods, x_i_,y_i_ denotes a point in space. In actual computation, when comparing multiple distributions, we calculates the median distance between all points across all distributions to obtain γ.

For structural validity, we calculated backbone bond lengths, bond angles, dihedral angles, and side-chain torsion angles to characterize the alignment of the local geometry of generated structures with MD trajectories. RMSD, RMSF, Rg and all local geometry metrics mentioned in this paper are calculated using MDTraj^42^.

### Fast Folding Evaluation Metrics

During the assessment of forward simulation LDT, to ensure that the assessment occurs at the same trajectory time step, we downsampled the MD test trajectory to 10,000 frames, the same as the training set. Additionally, since a single Fast Folding simulation already adequately samples various folding and unfolding events, we only used the first simulation of each protein during assessment. In each inference, DynaFold accepts an initial structure and predicted subsequent trajectories of 900 frames. The reverse diffusion parameters are consistent with ATLAS Evaluation Metrics.

For ensemble assessment, We firstly used MD trajectories to construct time-lagged independent components (TICA)^31,32^ mapping functions, and used the two slowest time-lagged independent components to describe the free energy surface of each trajectory for assessing the similarity between predicted equilibrium distribution and MD. Subsequently, we extracted the conformations corresponding to up to three local minima in the free energy surface and calculated backbone RMSD and RMSD of non-loop components to evaluate DynaFold’s ability to capture key conformations. In the actual evaluation process, since the low-free-energy center from the density estimation often lack real data points in many cases, we selected conformations within the vicinity of the local minima on the MD free-energy surface as alternative conformations. The pair with the lowest backbone RMSD was used as the matching result between model-predicted conformations and MD alternative conformations. Finally, We calculated the secondary structure distribution, residue contact frequency and solvent accessible surface area to reflect the folding pattern and folding trajectories predicted by DynaFold.

During the assessment of conformational transition LDT, we uniformly downsampled trajectories to 100,000 frames to capture conformational changes in greater detail than the forward simulation model. Subsequently, we extracted three distinct events from each protein: α-helix transformation into β-sheet, β-sheet folding, and α-helix unfolding. The MD sub-trajectories for these three events were obtained by identifying segments on the trajectories satisfying the following three conditions: 1) The distance between the start and end points on the TICA graph exceeded 0.2 times the sum of the x-axis range and y-axis range, ensuring low temporal correlation between the two conformations and requiring a sufficient trajectory length to describe the conformational transition. 2) Maximum changes in secondary structure composition, thereby covering the most complex conformational transition scenarios within the trajectory. 3) Up to 100 frames. In each inference, DynaFold accepts an initial and a final structure, and samples conformational transition trajectories totalling 10 or 50 frames. The reverse diffusion parameters are consistent with ATLAS Evaluation Metrics, except that the number of denoising steps has been reduced to 100.

The conformation transition assessment calculated three metrics: TICA space pathways, secondary structure changes, and backbone RMSD variations, so as to evaluate the conformational changes sampled by DynaFold within both the MD-fitted TICA space and three-dimensional space. The TICA space was fitted using downsampled trajectories from the first MD simulation of the corresponding fast-folding protein.

## Acknowledgement

This work was supported by Shanghai Municipal Science and Technology Major Project, partially by SJTU Kunpeng & Ascend Center of Excellence, the Center for HPC at Shanghai Jiao Tong University, and the National Key Research and Development Program of China (2025YFA0921000 and 2023YFF1205102), the Fundamental Research Funds for the Central Universities (YG2023LC03), and the National Natural Science Foundation of China (32171242).

## Data and Code Availability

Source code and model parameters are available at https://github.com/Zirui-Fan/DynaFold.

## Notes

**Notes** The authors declare that there is no conflict of interest.

### Competing Interest Statement

The authors have declared no competing interest.

### Summary of Updates

A structural ensemble prediction model has been incorporated, with corresponding modifications made to the figures within the article and supplementary file.

