## Supplementary Information for "DynaFold: A Latent Diffusion Based Generative Framework for Protein Dynamic Trajectory"

**Notes**

The authors declare that there is no conflict of interest.

### Supplementary algorithms

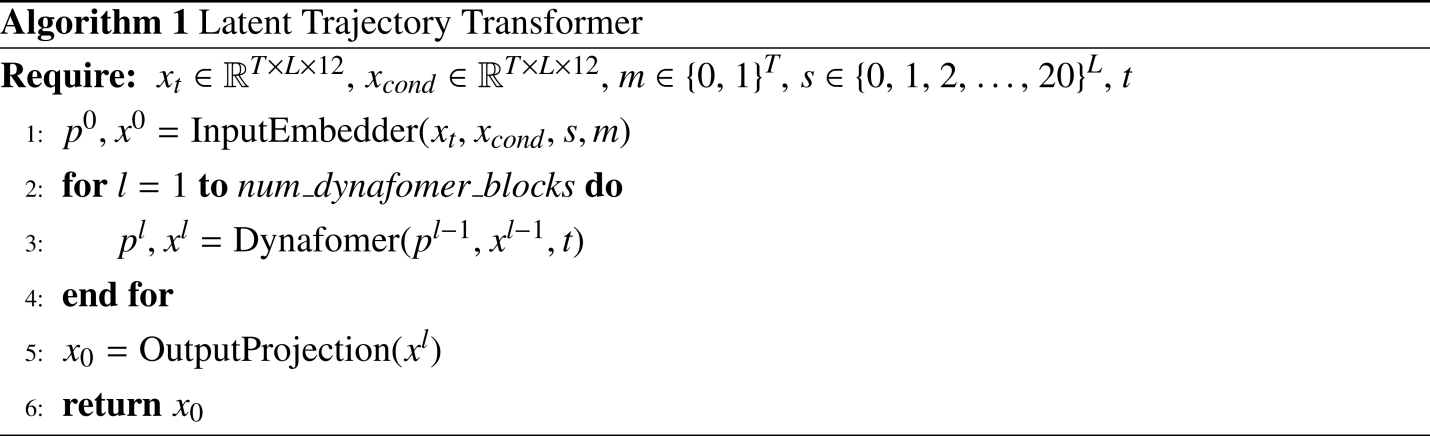

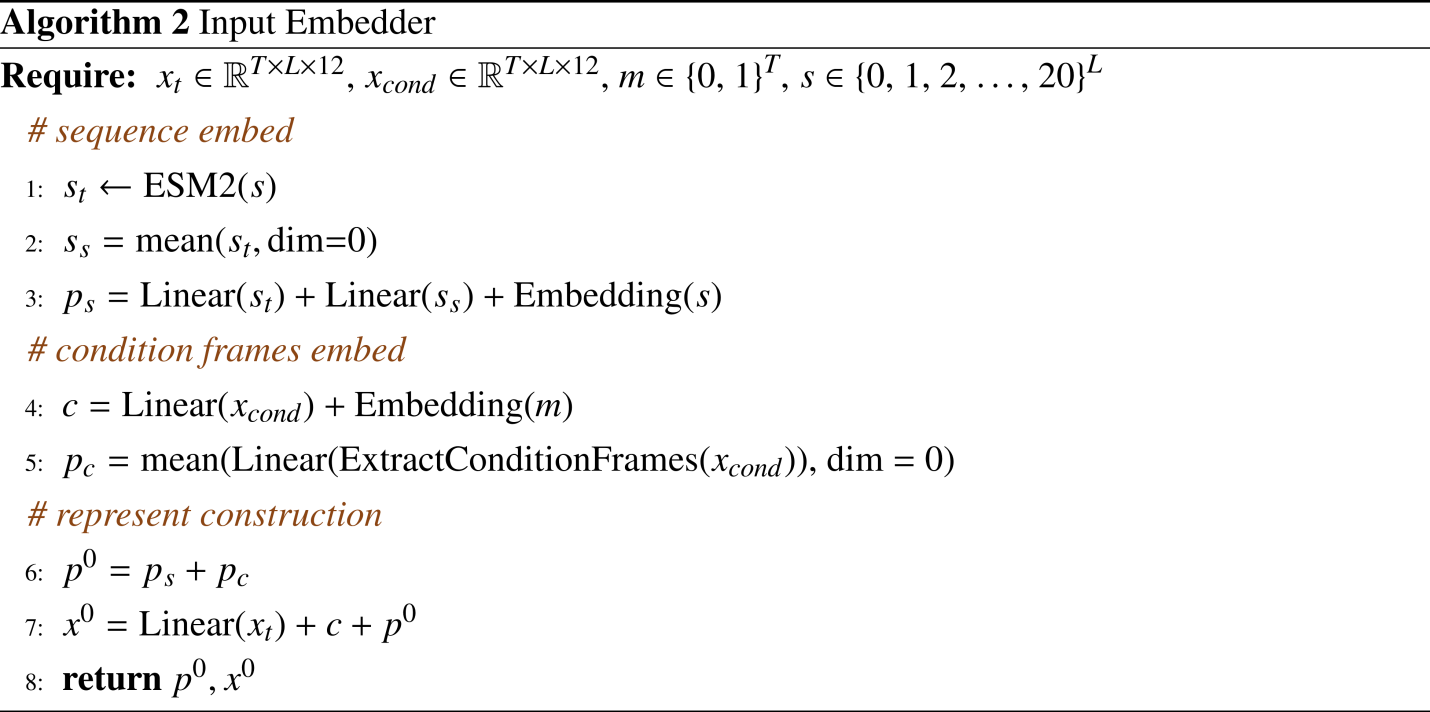

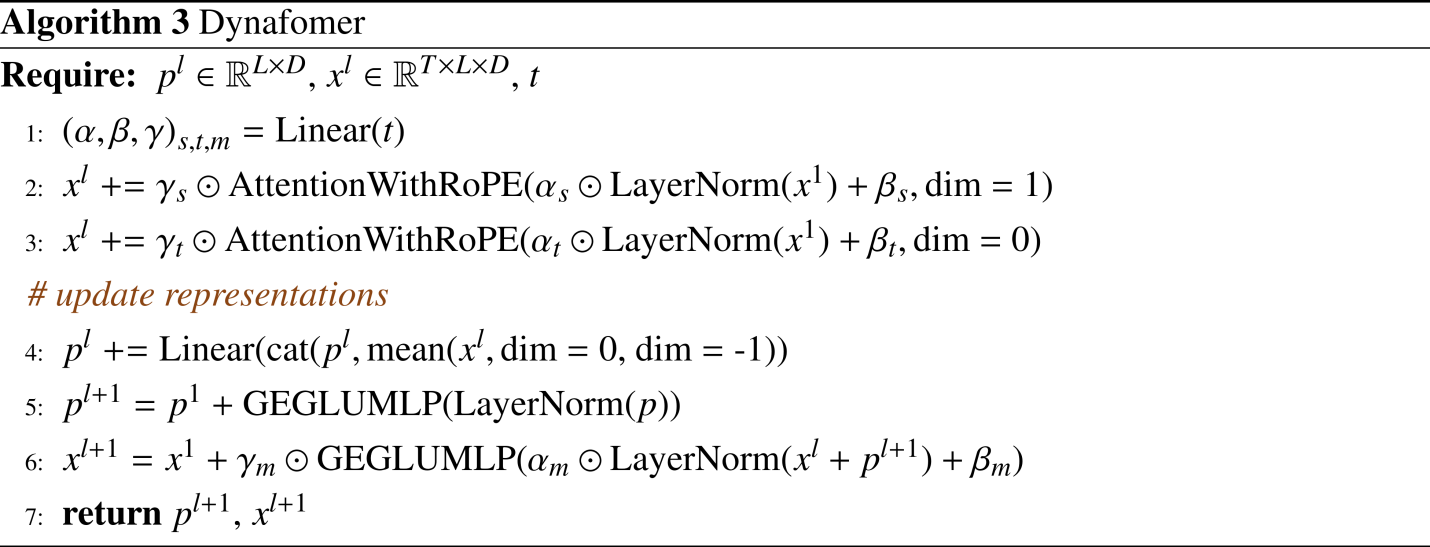

### Supplementary figures

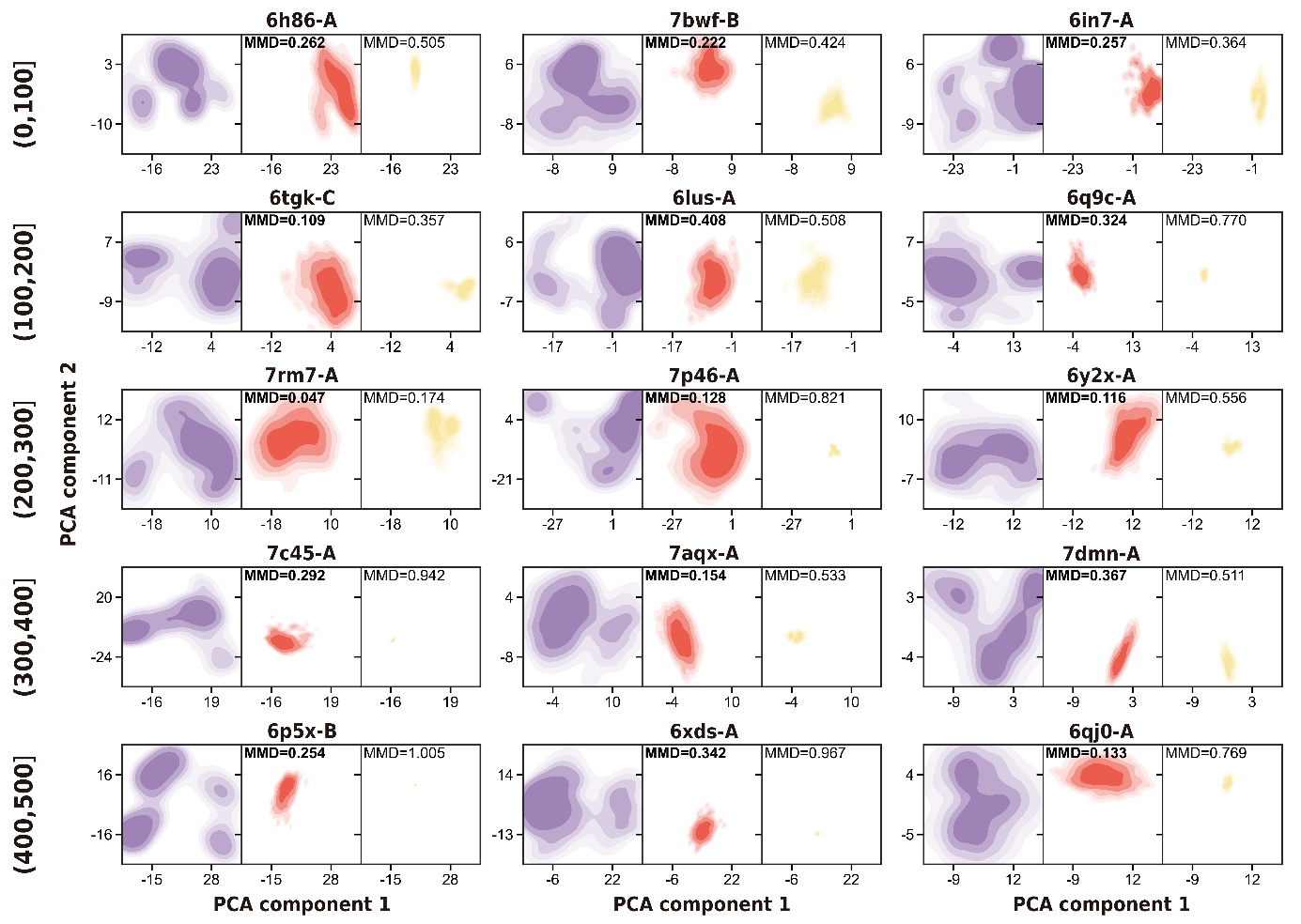

**Figure S1.** Distribution of MD predicted ensembles and different model predicted ensembles in PCA space.

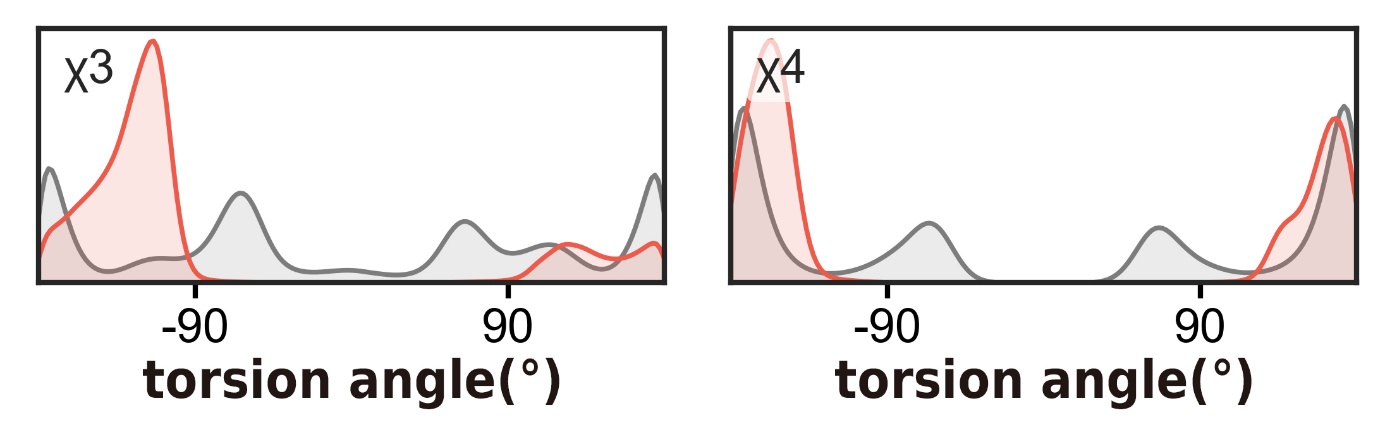

**Figure S2.** χ3-χ4 distributions of MD predicted ensembles and DynaFold predicted ensembles. DynaFold: red, MD: grey.

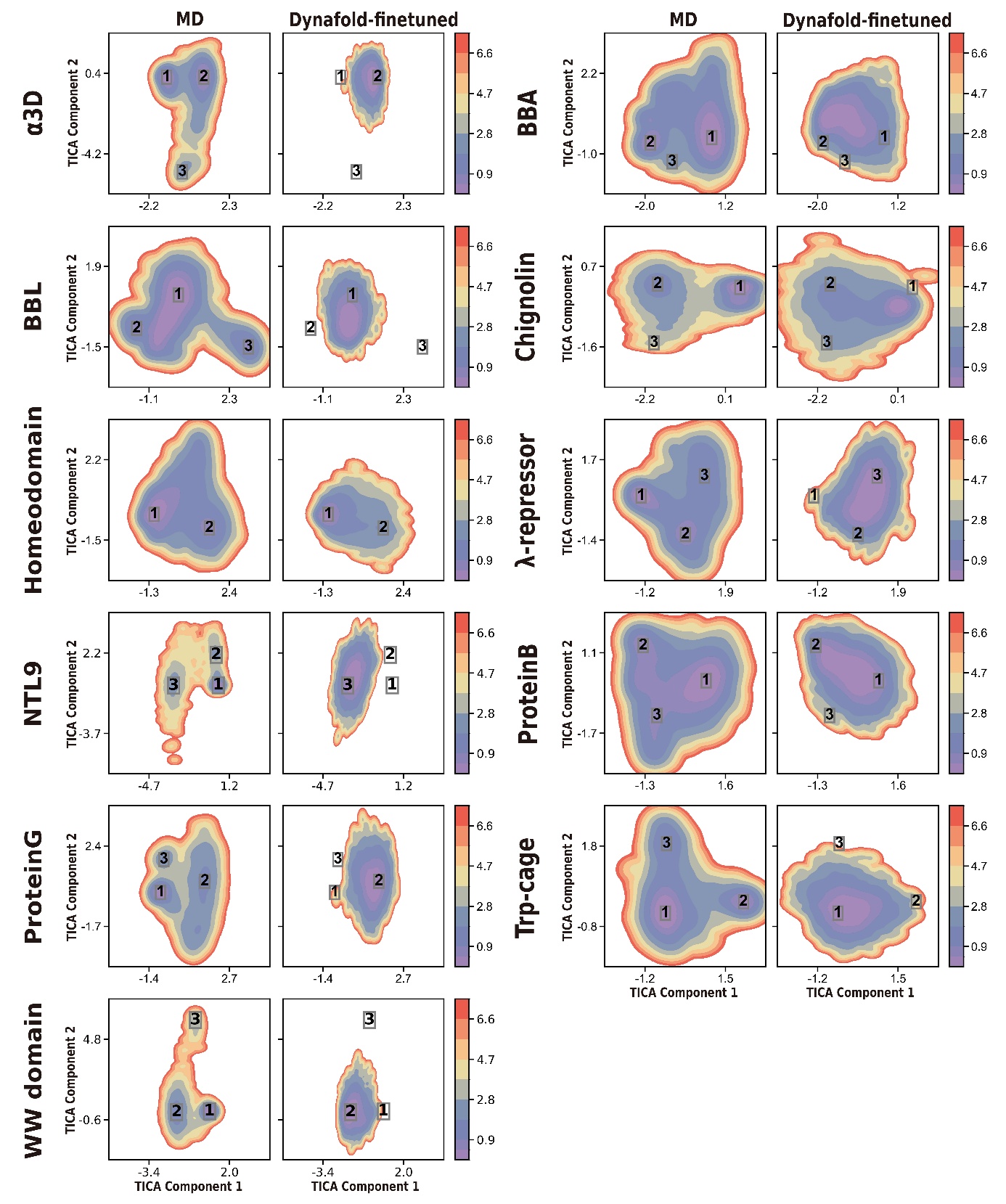

**Figure S3.** Free energy surface of MD ground truth and DynaFold's prediction in MD-fitted TICA space.

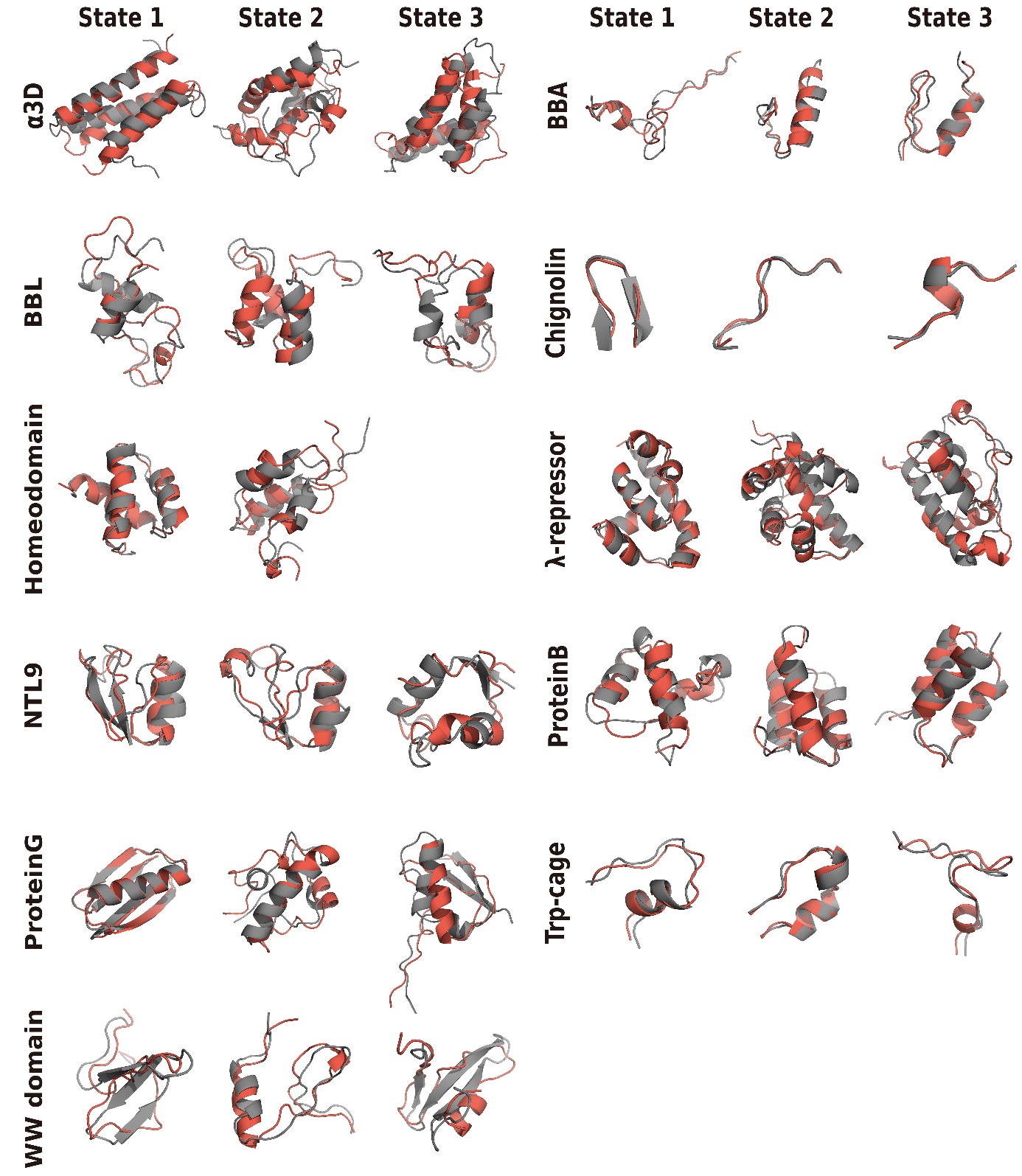

**Figure S4.** Low-free-energy conformations of MD ground truth and DynaFold's prediction.

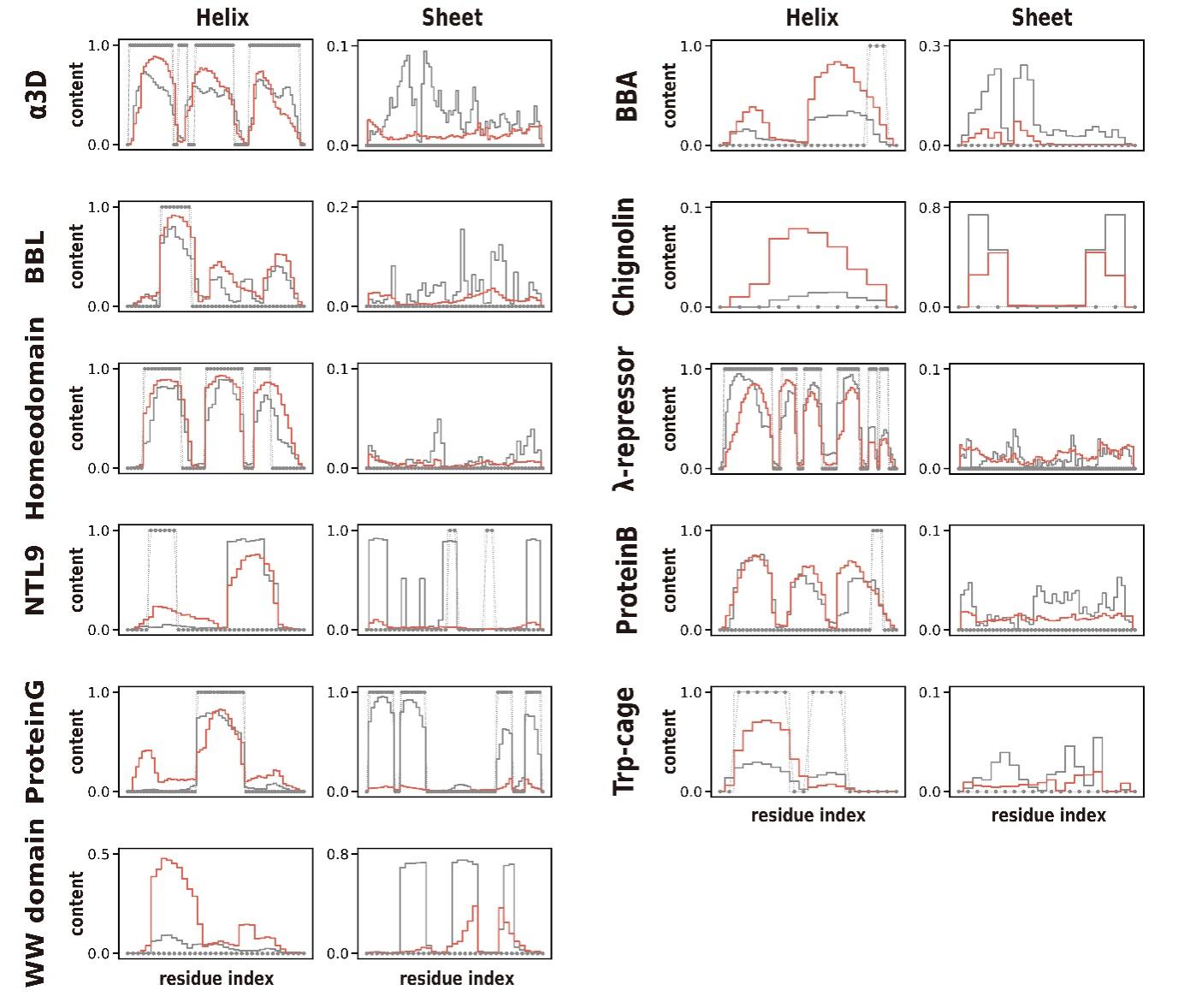

**Figure S5.** Secondary structure frequency of MD ground truth and DynaFold's prediction.

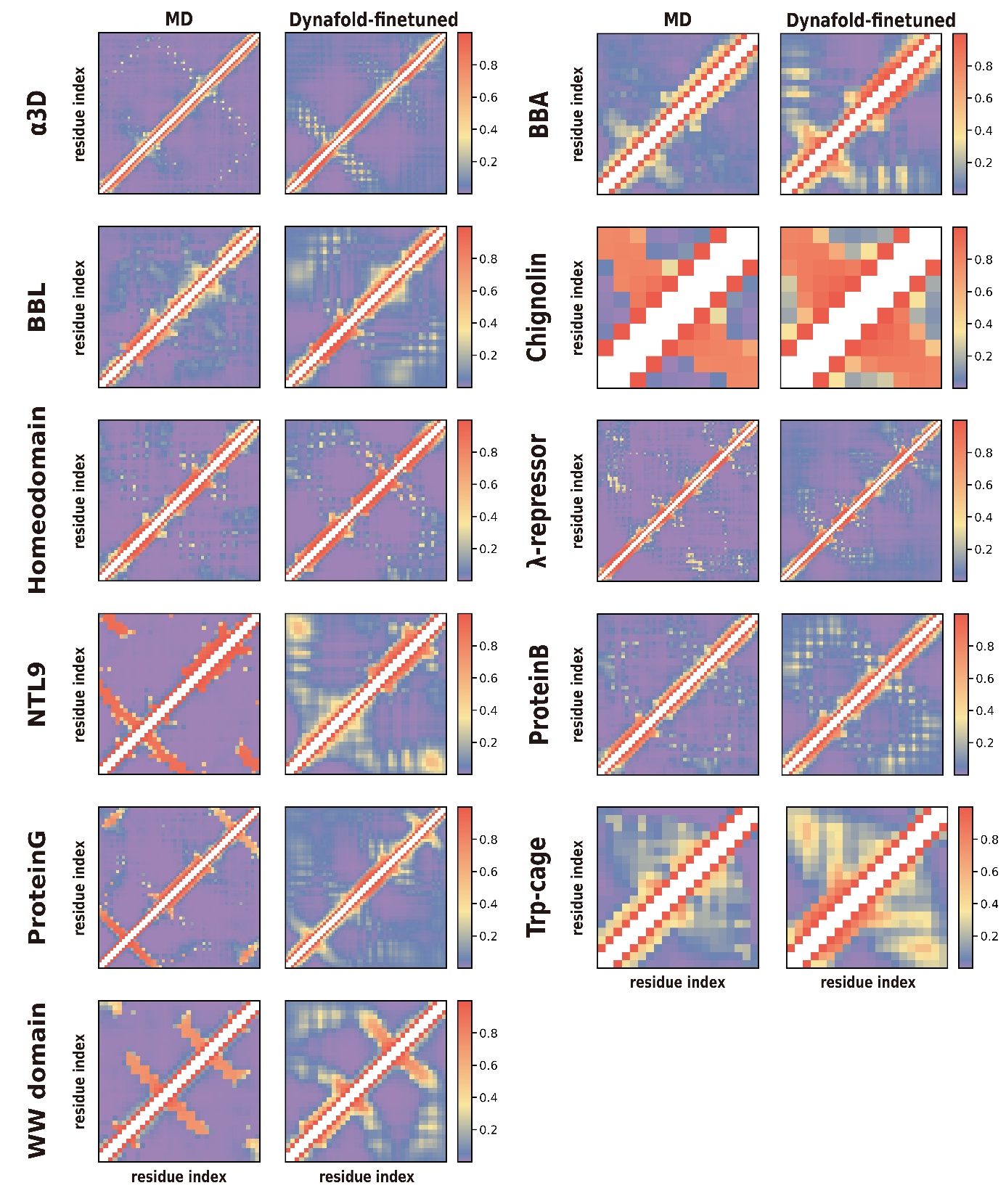

**Figure S6.** Residue contact frequency of MD ground truth and DynaFold's prediction.

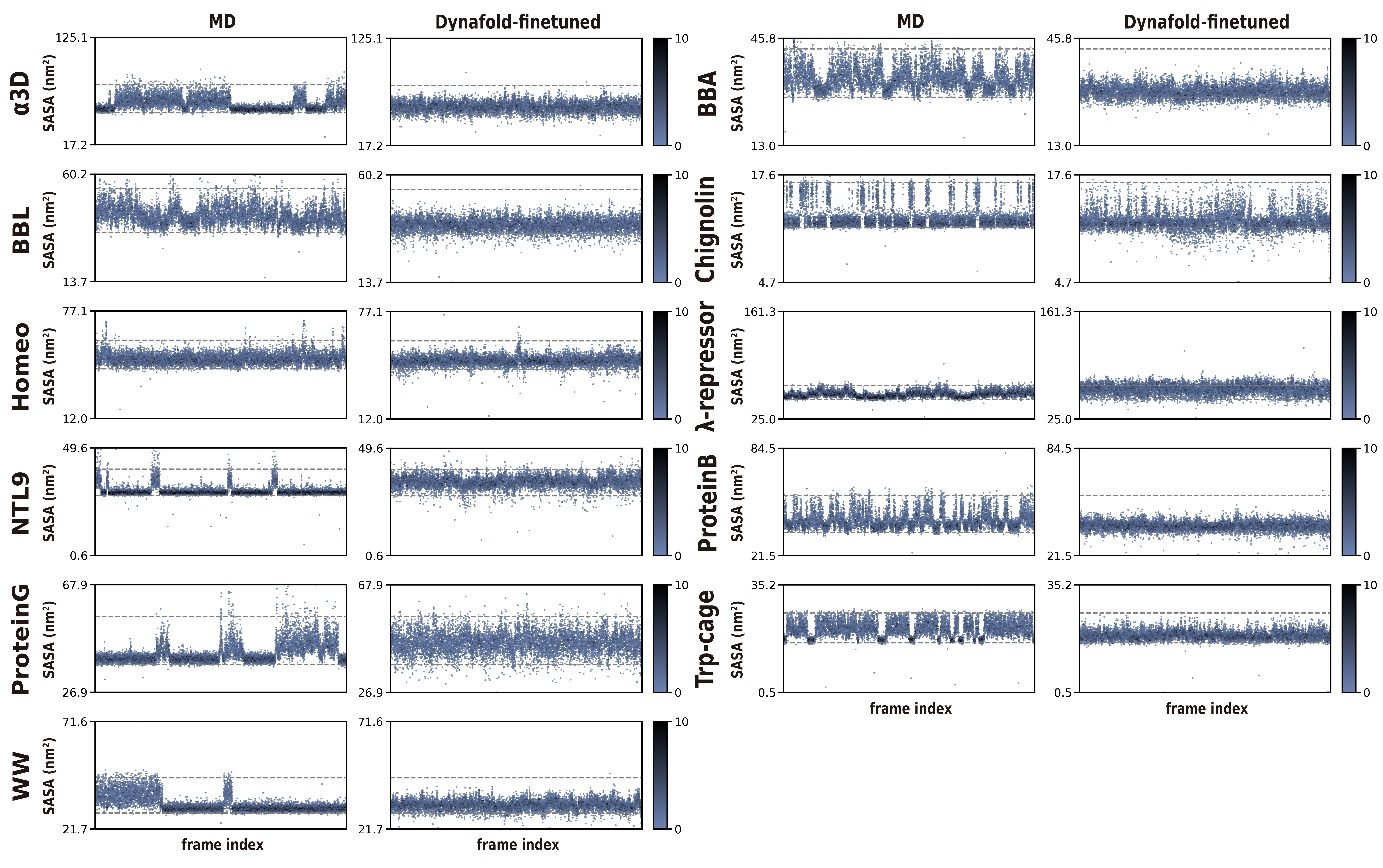
**Figure S7.** Solvent accessible surface areas (SASA) of MD ground truth and DynaFold's prediction in MD-fitted TICA space.

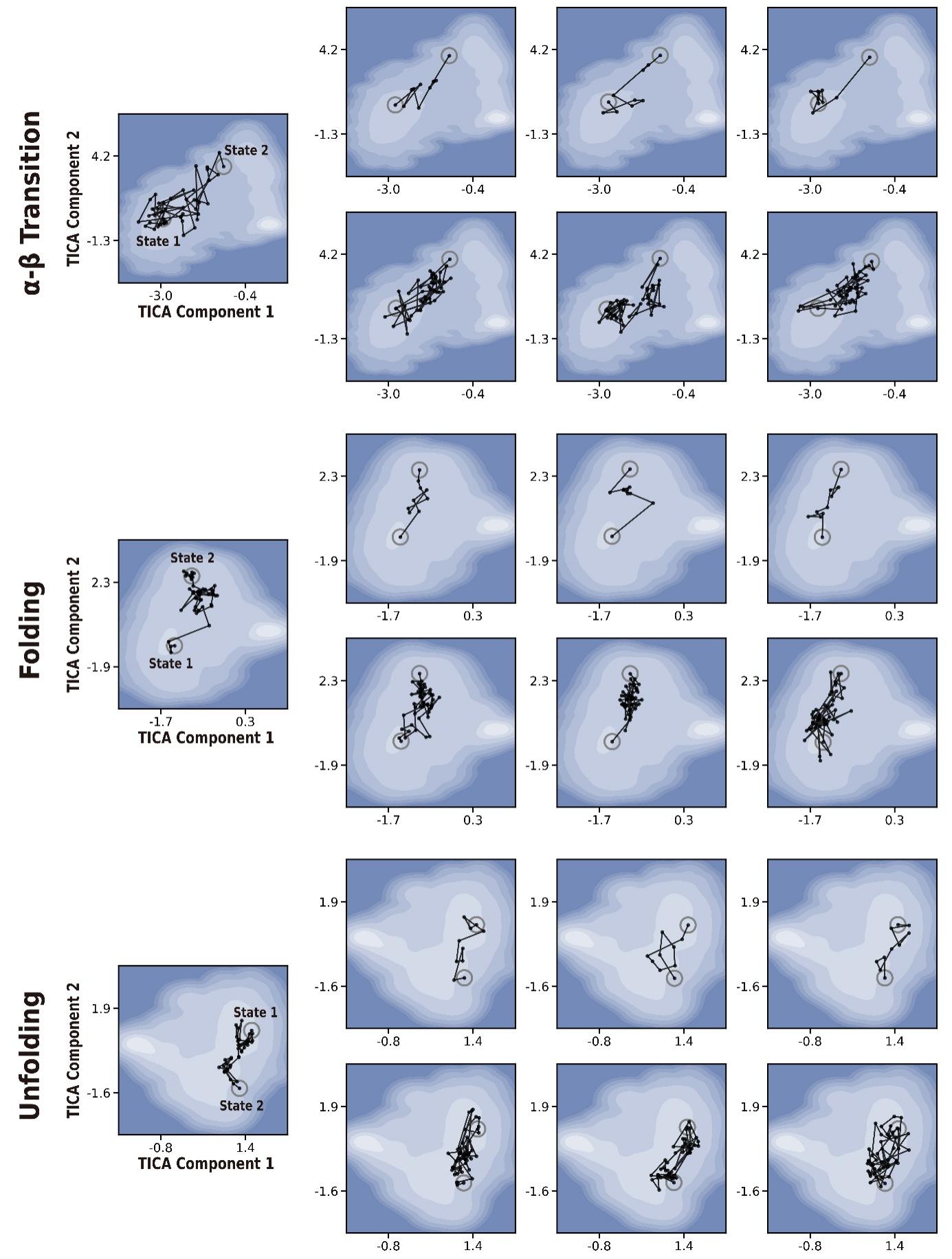
**Figure S8.** Various conformational transition pathways predicted by DynaFold.

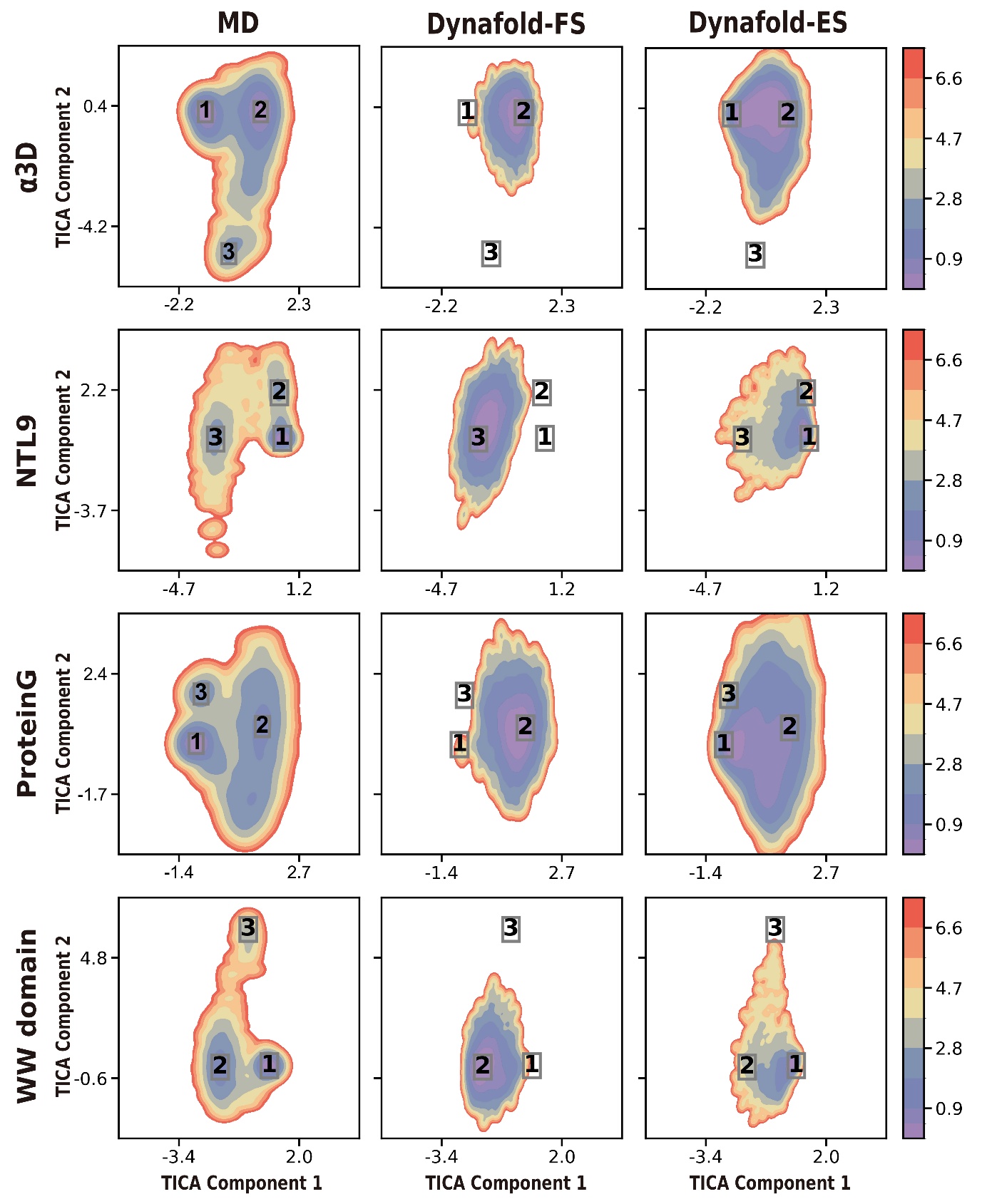

**Figure S9.**  Free energy surface of MD ground truth, DynaFold-FS and DynaFold-ES prediction in MD-fitted TICA space.

### Supplementary tables

**Table S1.** Top three ATLAS test set proteins with the highest average RMSF (Å) within five length intervals.

| **Bin** | **Protein ID** | **Length** | **Avg RMSF** |
| --- | --- | --- | --- |
| (0,100] | 6h86-A | 88 | 0.579 |
|  | 7bwf-B | 92 | 0.566 |
|  | 6in7-A | 81 | 0.446 |
| (100,200] | 6tgk-C | 105 | 0.441 |
|  | 6lus-A | 133 | 0.277 |
|  | 6q9c-A | 155 | 0.242 |
| (200,300] | 7rm7-A | 228 | 0.381 |
|  | 7p46-A | 282 | 0.331 |
|  | 6y2x-A | 235 | 0.317 |
| (300,400] | 7c45-A | 302 | 0.363 |
|  | 7aqx-A | 364 | 0.205 |
|  | 7dmn-A | 377 | 0.122 |
| (400,500] | 6p5x-B | 457 | 0.445 |
|  | 6xds-A | 492 | 0.311 |
|  | 6qj0-A | 412 | 0.210 |

**Table S2.** Maximum Mean Discrepancy between MD predicted ensembles and different model predicted ensembles in PCA space and Rg-RMSD space.

| **Protein ID** | **DynaFold** | **MDGen** |
| --- | --- | --- |
| 6h86-A | **0.262** | 0.505 |
| 7bwf-B | **0.257** | 0.364 |
| 6in7-A | **0.408** | 0.508 |
| 6tgk-C | **0.254** | 1.006 |
| 6lus-A | **0.324** | 0.77 |
| 6q9c-A | **0.133** | 0.769 |
| 7rm7-A | **0.109** | 0.357 |
| 7p46-A | **0.342** | 0.967 |
| 6y2x-A | **0.116** | 0.556 |
| 7c45-A | **0.154** | 0.533 |
| 7aqx-A | **0.222** | 0.424 |
| 7dmn-A | **0.292** | 0.942 |
| 6p5x-B | **0.367** | 0.511 |
| 6xds-A | **0.128** | 0.822 |
| 6qj0-A | **0.047** | 0.174 |
| Mean | **0.228** | 0.614 |
| Minimum count | **15** | 0 |

**Table S3.** Backbone RMSD (Å) between MD low-free-energy conformations and the most similar conformations predicted by DynaFold.

| **Protein** | **1** | **2** | **3** | **Mean** |
| --- | --- | --- | --- | --- |
| Alpha3D370K | 4.27 | 5.07 | 6.07 | 5.13 |
| BBA325K | 2.76 | 1.62 | 1.33 | 1.90 |
| BBL298K | 3.84 | 3.35 | 3.78 | 3.65 |
| Chignolin340K | 0.50 | 0.93 | 0.89 | 0.77 |
| Homedomain360K | 1.66 | 3.37 | — | 2.52 |
| Lambdarepressor350K | 1.12 | 3.82 | 4.47 | 3.14 |
| NTL9355K | 2.53 | 2.35 | 2.92 | 2.60 |
| ProteinB340K | 3.45 | 1.71 | 1.99 | 2.38 |
| ProteinG350K | 0.76 | 3.65 | 4.04 | 2.81 |
| Trpcage290K | 1.65 | 0.67 | 2.09 | 1.47 |
| WW360K | 2.85 | 2.80 | 3.50 | 3.05 |
| **Mean** | — | — | — | 2.68 |

**Table S4.** Secondary structure RMSD (Å) between MD low-free-energy conformations and the most similar conformations predicted by DynaFold. Calculated only on structures with a secondary structure content exceeding 40%.

| **Protein** | **1** | **2** | **3** | **Mean** |
| --- | --- | --- | --- | --- |
| Alpha3D370K | 1.55 | 3.08 | 4.09 | 2.90 |
| BBA325K | — | 0.95 | 0.55 | 0.75 |
| BBL298K | — | 2.08 | — | 2.08 |
| Chignolin340K | 0.43 | — | 0.30 | 0.36 |
| Homedomain360K | 1.24 | 1.89 | — | 1.57 |
| Lambdarepressor350K | 0.96 | 3.01 | 2.31 | 2.09 |
| NTL9355K | 1.31 | 1.49 | 2.45 | 1.75 |
| ProteinB340K | 3.15 | 1.21 | 1.50 | 1.95 |
| ProteinG350K | 0.71 | 1.22 | 2.90 | 1.61 |
| Trpcage290K | — | 0.32 | — | 0.32 |
| WW360K | — | — | — | — |
| **Mean** | — | — | — | 1.68 |

**Table S5.** Maximum Mean Discrepancy between MD predicted ensembles and DynaFold predicted ensembles in 100000-frame MD-fitted TICA space.

| **Protein** | **DynaFold-ES** |
| --- | --- |
| Alpha3D370K | 0.037 |
| NTL9355K | 0.237 |
| ProteinG350K | 0.083 |
| WW360K | 0.110 |
| **Mean** | 0.117 |

**Table S6.** Maximum Mean Discrepancy between MD predicted ensembles and DynaFold predicted ensembles in 10000-frame MD-fitted TICA space.

| **Protein** | **DynaFold-FS** | **DynaFold-ES** |
| --- | --- | --- |
| Alpha3D370K | 0.217 | 0.127 |
| NTL9355K | 1.077 | 0.333 |
| ProteinG350K | 0.553 | 0.155 |
| WW360K | 0.626 | 0.142 |
| **Mean** | 0.618 | 0.189 |
